# Predicting T Cell Receptor Functionality against Mutant Epitopes

**DOI:** 10.1101/2023.05.10.540189

**Authors:** Emilio Dorigatti, Felix Drost, Adrian Straub, Philipp Hilgendorf, Karolin I. Wagner, Bernd Bischl, Dirk H. Busch, Kilian Schober, Benjamin Schubert

**Affiliations:** Department of Statistics, Ludwig Maximilian Universität, Munich, Germany; Institute of Computational Biology, Helmholtz Zentrum München — German Research Center for Environmental Health, Neuherberg, Germany; Munich Center for Machine Learning (MCML); School of Life Sciences Weihenstephan, Technical University of Munich, Munich, Germany; Institute for Medical Microbiology, Immunology, and Hygiene, Technical University of Munich, Munich, Germany; Mikrobiologisches Institut–Klinische Mikrobiologie, Immunologie und Hygiene, Universitätsklinikum Erlangen, Friedrich-Alexander-Universität Erlangen-Nürnberg, Erlangen, Germany; German Center for Infection Research, Deutschen Zentrum für Infektionsforschung (DZIF), Partner Site Munich, Munich, Germany; Medical Immunology Campus Erlangen, Friedrich-Alexander-Universität (FAU) Erlangen-Nürnberg, Schlossplatz 1, Erlangen, Germany; Department of Mathematics, Technical University of Munich, Garching bei München, Germany

## Abstract

Cancer cells or pathogens can escape recognition by T cell receptors (TCRs) through mutations of immunogenic epitopes. TCR cross-reactivity, i.e., recognition of multiple epitopes with sequence similarities, can be a factor to counteract such mutational escape. However, cross-reactivity of cell-based immunotherapies may also cause severe side effects when self-antigens are targeted. Therefore, the ability to predict the effect of mutations in the epitope sequence on T cell functionality *in silico* would greatly benefit the safety and effectiveness of newly-developed immunotherapies and vaccines. We here present “Predicting T cell Epitope-specific Activation against Mutant versions” (P-TEAM), a Random Forest-based model which predicts the effect of point mutations of an epitope on T cell functionality. We first trained and tested P-TEAM on a comprehensive dataset of 36 unique murine TCRs in response to systematic single-amino acid mutations of their target epitope (representing 5.472 unique TCR-epitope interactions). The model was able to classify T cell reactivities, corresponding to *in vivo* recruitment of T cells, and quantitatively predict T cell functionalities for unobserved single-point mutated altered peptide ligands (APLs), or even unseen TCRs, with consistently high performance. Further, we present an active learning framework to guide experimental design for assessing TCR functionality against novel epitopes, minimizing primary data acquisition costs. Finally, we applied P-TEAM to a novel dataset of 7 human TCRs reactive to the tumor neoantigen VPSVWRSSL. We observed a similarly robust performance for these human TCRs as for the murine TCRs recognizing SIINFEKL, thus providing evidence that our approach is applicable to therapeutically relevant TCRs as well as across species. Overall, P-TEAM provides an effective computational tool to study T cell responses against mutated epitopes.

## Introduction

The TCR-mediated recognition of pathogen- or tumor-derived epitopes by T cells plays an essential role in the adaptive immune response. These epitopes are bound by the Major Histocompatibility Complex (MHC), and interact with the complementarity-determining regions (CDRs) of the T cell receptor (TCR). T cells whose TCR recognizes the epitope with sufficient affinity are activated and undergo clonal expansion and differentiation to form an immune response. The exchange of a single amino acid in the epitope may severely alter TCR binding behaviour [1]. Further, undetected cross-reactivity towards healthy cells may cause severe damage when developing T cell-based immunotherapies against neo-epitopes from tumor cells and must therefore be avoided [2]. In order to ensure the safety and effectiveness of immunotherapies and vaccines, it is therefore fundamental to understand the effects of epitope mutations on TCR recognition.

Computationally predicting binding between TCR and epitope remains challenging due to the immense sequence diversity: it has been estimated that there exist more than 10^20^ possible TCRs in nature and that every human harbors at least 10^7^ different TCRs at any given time [3]. Clustering and distance algorithms are able to estimate TCR specificity by comparing multiple receptors, mainly based on their CDR3*β* sequence [4–6]. Additionally, publicly available, curated datasets with paired information on TCRs and their recognized epitopes allowed the creation of a variety of machine learning methods to predict TCR-epitope binding [6–11]. However, these data are not collected in a standardized manner such as deep mutational epitope scans: as of April 2023, for example, only 17 of the 152 single-amino acid mutated peptides for the model epitope SIINFEKL are provided in the IEDB [12] and none in the VDJdb [13]. Therefore, current methods trained on such data are likely to fail when predicting the change in T cell activation introduced by most point mutations. Additionally, these databases typically simplify the TCR-epitope interaction to a binary event of binding or non-binding, even though epitopes activate T cells to various degrees, resulting in continuous changes in the phenotype and abundance of T cell populations during an immune response [14]. Finally, it is usually not defined if and which binding levels correspond to *in vivo* recruitment and functionality of T cell clones. Predictors trained on binary TCR binding labels, therefore, miss most of these nuanced functional changes which can be introduced through single amino acid mutations.

The dataset from a deep mutational scan introduced by Straub et al. [15] tackles both of these shortcomings by measuring the effect of all single-point mutations of the well-characterized murine epitope SIINFEKL on TCR functional reactivity, and simultaneously determining T cell reactivity levels that correspond to actual recruitment and clonal expansion *in vivo* after pathogen infection (Figure 1a). In this work, we leverage this dataset to introduce P-TEAM, a Random Forest model trained to predict how the T cell reactivity is affected by single amino acid altered peptide ligands (APLs, Figure 1b). The model can either learn from a TCR’s reactivities towards a subset of APLs to predict the effect of the remaining mutations, or generalize across a fully characterized TCR repertoire to novel TCRs. In contrast to most previous predictors, P-TEAM’s model does not only classify TCR-epitope pairs as binding or non-binding but is also able to estimate a continuous activation score reflective of TCR reactivity. Additionally, we embed P-TEAM into an active learning framework for experimental design to reduce the amount of training data required to obtain reliable predictions for novel epitopes. Finally, we apply P-TEAM on TCRs with therapeutic potential reactive to the human cancer neo-antigen VPSVWRSSL, showing high performance in predicting potential cross-reactive epitopes for T cell-based immunotherapies.

**Figure 1.**
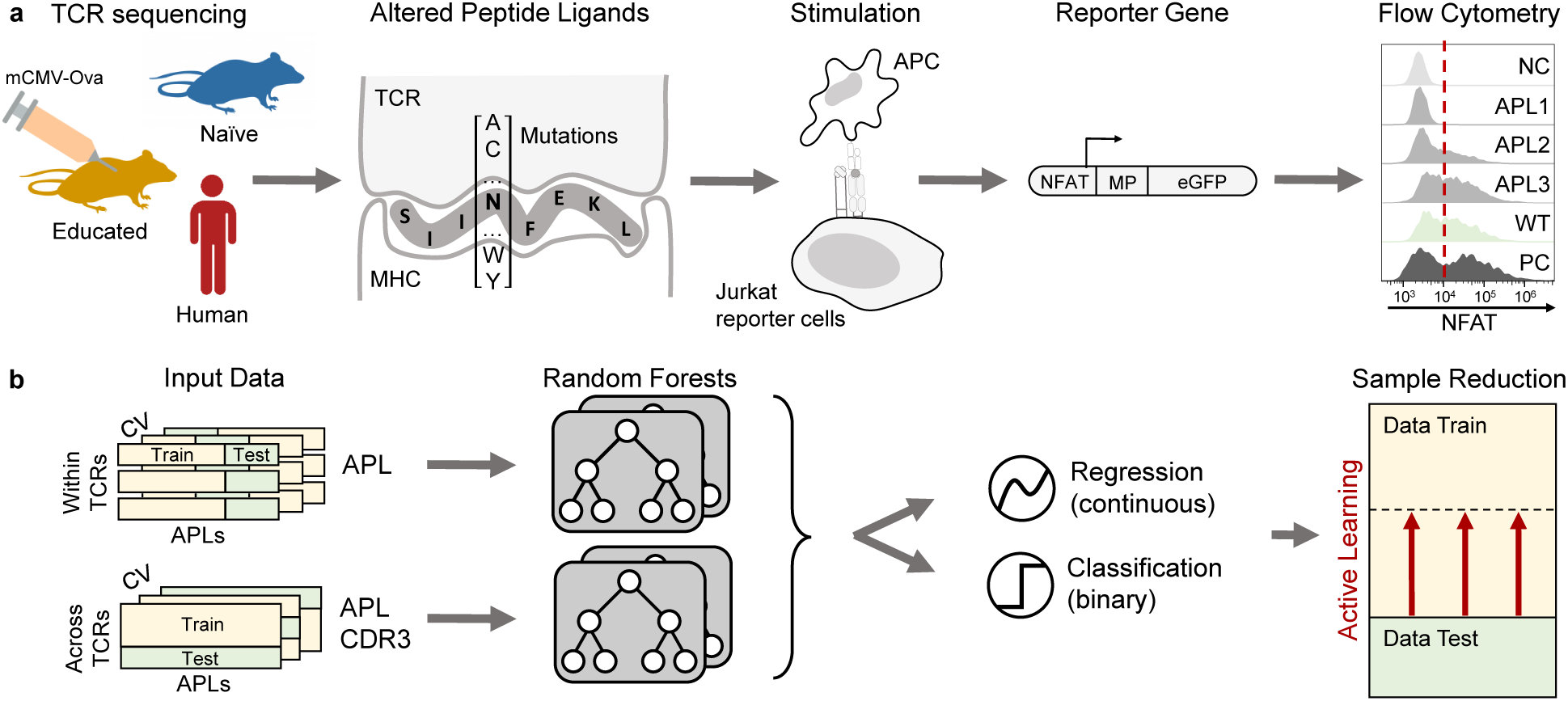
Overview of P-TEAM for predicting T cell activation by mutational epitopes. **a**, Data acquisition: SIINFEKL-reactive TCRs were isolated based on H2k^b^-SIINFEKL multimers from mice previously exposed or unexposed to a murine Cytomegalovirus strain expressing SIINFEKL (mCMV-Ova). An additional dataset consists of six previously identified human TCRs reactive to the tumor epitope VPSVWRSSL. Jurkat triple parameter reporter cells (JTPR) expressing a single TCR each were stimulated with altered peptide ligands (APLs) derived from single amino acid mutations of the cognate epitopes. An activation score of each TCR towards each APL, a negative (NC) and positive control (PC), and the wild-type epitope (WT) is determined based on NFAT expression measured by flow cytometry after stimulation. **b**, Random Forests predict the continuous activation score as a regression task, or the binary TCR recognition related to *in vivo* recruitment as a classification task, for all APLs within a TCR, as well as for an unseen TCR. The amount of training data can be reduced by efficient sampling using active learning techniques.

## Results

### Comprehensive quantification of T cell reactivity towards single-point mutations

In order to develop a model for predicting T cell functionality for mutational epitopes, we utilized the novel dataset described in [15], which contains functional reactivity information of 36 murine TCR sequences towards all single-mutation based APLs of the model epitope SIINFEKL (murine dataset). The TCRs were either isolated from the naïve repertoire of SIINFEKL-binding T cells (*n* = 20) of an unexposed mouse [15] or the memory repertoire (*n* = 15) of murine Cytomegalovirus (mCMV)-SIINFEKL exposed mice [16]. For the sake of brevity, these subsets of the murine dataset are further referred to as “naïve repertoire” and “educated repertoire,” respectively. The dataset also included the well-studied SIINFEKL-reactive TCR OT-I as a reference control (*n* = 1).

For each TCR, an activation score representing the fraction of activated T cells (Supplementary Figure 1a) was experimentally determined for the wild-type epitope as well as all of its 152 APLs (at each of the 8 peptide positions one of the 19 other amino acids was used at a time; 8×19=152; Methods Data Collection). In total, we studied 5,472 (36 TCRs x 152 APLs) unique murine TCR-pMHC interactions. The scores were scaled linearly to normalize the data (Figure 2a) across the different TCRs. All TCRs were tested in one experiment against the wild-type epitope. In subsequent experiments, each TCR at a time was tested against all APLs. The normalization factor was then derived as the fraction of the activation scores towards the wild-type epitope between an experiment including all TCRs simultaneously and the individual runs in which one TCR at a time was investigated. A TCR was marked as reactive to a peptide when the activation score exceeded 46.9%, which was identified as a threshold for effective recruitment and clonal expansion *in vivo* upon infection with Listeria monocytogenes expressing the SIINFEKL epitope [15].

**Figure 2.**
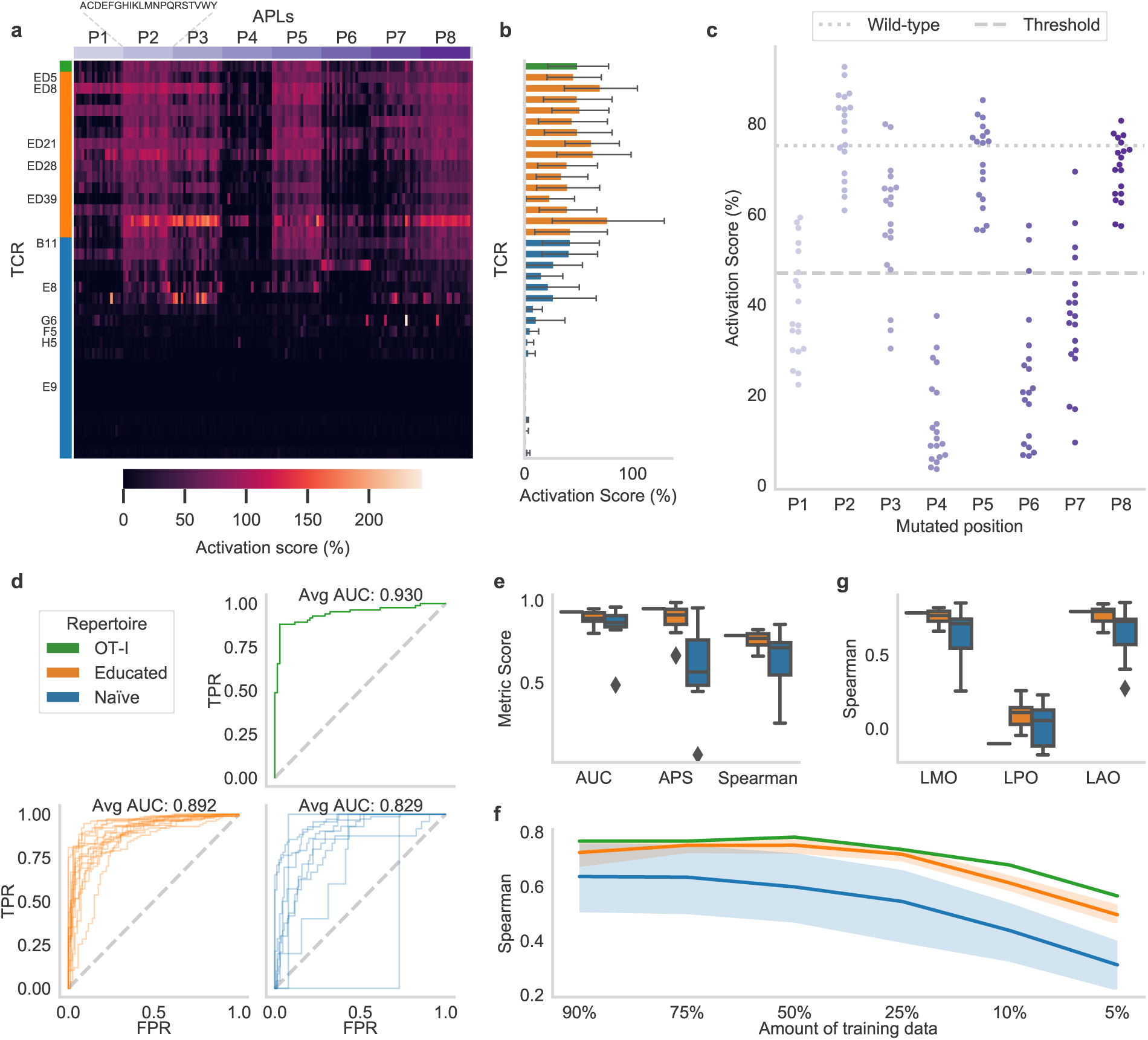
Predicting APL mutations within the reactivity landscape of a given TCR. **a**, The normalized activation scores show a large variety of murine T cell activation in the deep mutational scan, in which each of the eight epitope position (P1-P8) was exchanged by the other 19 amino acids in turn. **b**, The activation scores averaged for all APLs of one TCR indicate high- and low-affinity TCRs. **c**, The epitope position on which the mutation occurs strongly influences the activation per APL (*n* = 19) averaged over all TCRs (*n* = 36). The threshold value represents the boundary between binding and non-binding and wild-type indicates the activation scores of the base epitope SIINFEKL. **d**, The Receiver operating characteristic (ROC) curves for the different TCR repertoires indicate the True Positive Rate (TPR) against the False Positive Rate (FPR) at all prediction values as thresholds. **e**, Different evaluation metrics for regression (Spearman) and classification models (APS: Average Precision Score, AUC: Area Under the ROC Curve). **f**, Spearman correlation when a smaller amount of training data is used (average over ten repetitions with random subsets for each TCR). The boundaries of the mean line indicate the 95% confidence interval. **g**, Spearman correlation obtained when trained on different subsets of the data. LMO: leave-mutation-out, LPO: leave-position-out, and LAO: leave-amino-acid-out. The box plot indicates the data quartiles while the whiskers extend to the extreme values excluding outliers. The median is indicated as a horizontal line.

Based on these experiments, the TCRs showed a broad range of functional reactivity levels (Figure 2b, Supplementary Figure 1b). Even though the TCRs were identified through binding toward H2k^b^-SIINFEKL multimers, 11 TCRs from the naïve repertoire did not show any relevant reactivity towards the wild-type epitope or any APL upon transgenic re-expression in Jurkat triple parameter reporter cells (JTPRs). Of these, 8 TCRs without reactivity towards any APL were excluded from all further experiments. 11 receptors were reactive with more than 50% of the peptides, with a maximum of 122 recognized APLs in one case. The model TCR OT-I recognized 94 APLs in addition to the wild-type sequence SIINFEKL. Overall, the change in activation score is highly dependent on the position of the mutation (Figure 2c, Supplementary Figure 1c). Exchanges at epitope positions 4 and 6 heavily decreased T cell activation, while positions 2, 5, and 8 expressed high robustness.

In summary, our dataset contains murine TCRs derived from the educated and naïve repertoire, with experimentally determined reactivities against all possible APLs. The TCRs from the educated repertoire all showed reactivity against SIINFEKL and large numbers of its APLs, whereas TCRs from the naïve repertoire showed overall less, and more variable, reactivities against the wild-type and mutant epitope versions.

### P-TEAM predicts the effect of epitope point mutations on individual TCRs

We applied P-TEAM to this datasets to predict the effect of the various mutations of the epitope SIINFEKL for each TCR individually. Our approach is based on a Random Forest estimator, which receives physiochemical representations of amino acids [17] describing the wild-type epitope sequence and the APL sequences. The properties of each amino acid are described by five so-called Atchley Factors which summarize polarity, secondary structure, molecular size, amino acid composition in proteins, and electrostatic charge. To learn the effect of mutations in a data-driven manner, the model uses this numeric representation as input features which provide the characteristics of the APLs in a machine-comprehensible way.

We first investigated whether it is possible to predict recognition of APLs by TCRs, defined by the reactivity levels related to *in vivo* recruitment, using a binary classification model. To this end, we trained and evaluated the model in a leave-mutation-out cross-validation scheme using all data related to an individual TCR: The Random Forest was trained on 151 out of 152 APLs of one TCR and predicted the probability of activation of the left-out APL of this TCR. This process was repeated by excluding each APL in turn from the training set, and separately for each TCR. To evaluate the classification models, we reported the Area Under the Receiver Operating Characteristic Curve (AUC), which summarizes the True Positive Rate (TPR, recall, fraction of correct positive predictions over positive samples) and False Positive Rate (FPR, incorrect positive predictions over negative samples) at all possible classification thresholds. We further provide the Average Precision Score (APS), which indicates the area under the curve between precision (PPV, fraction of positive samples among those predicted as positive) and recall (see TPR). The AUC can be interpreted as the probability that a positive sample is scored higher than a negative sample, and is thus a natural and common metric used in binary classification tasks, while the APS is robust towards imbalanced data, thus providing a reliable performance measure even when most events are negative, as is the case for some TCRs in the dataset [18].

The performance (Supplementary Table 1) on the educated repertoire was consistently high with a maximal AUC of 94.9% and APS of 98.7% for the best predictable TCR ED21, and a mean of 89.2% and 88.3% among all TCRs, respectively (Figure 2d-e, Supplementary Figure 2a). Only two receptors in the educated repertoire had an APS below 80%: TCR ED28 (66.8%) and TCR ED39 (66.2%). The model for the reference TCR OT-I showed a similarly high performance as the educated repertoire (AUC: 93.0%, APS: 94.9%). The predictive performance decreased for the naïve repertoire with a median AUC of 82.9% and APS of 58.1%. This was expected given the more variable and overall lower reactivities against the wild-type epitope and APLs in this repertoire compared to the educated repertoire (Figure 2a-b). Consistent with this, the naïve repertoire showed higher prediction variability between the different TCRs reaching up to an AUC of 95.9% for TCR B11. The worst-performing TCRs achieved only an APS of 6.2% and AUC of 48.6% for TCR G6 which expressed reactivity to only five APLs. However, the second worst TCR E8 follows after a great leap in performance of 44.7% and 82.1%, respectively (Figure 2d-e, Supplementary Figure 2a) indicating that the TCRs of the naïve repertoire were in fact predictable when they were reactive to a greater amount of APLs.

Moving beyond binary classification, we also predicted the continuous functional reactivity of T cells in a regression setting for a given APL in the same leave-mutation-out validation scheme separated by TCR. In this setting, we evaluated the models by Spearman’s rank correlation to eliminate possible differences in activation scores among different receptors due to normalization artifacts (Supplementary Table 1). The regression models showed similar variability as the classification models. However, the gap in median performance between the educated and naïve repertoires was considerably greater: 13.8 percentage points for Spearman compared to 6.3 percentage points for AUC (Figure 2e). Based on these results, we conclude that the TCR-epitope interaction cannot only be predicted as a binary recognition event but as a fine-grained continuous reactivity landscape when the respective annotation is provided.

### Only 25% of random mutations are needed to learn a general model

We showed that P-TEAM can predict T cell reactivity levels, corresponding to *in vivo* recruitment, of a single mutation when trained on the remaining APLs. However, experimentally determining the activation of TCRs for the majority, if not all, of possible APLs comes with extensive labor, time, and cost expenses. Therefore, we analyzed the minimum amount of APLs needed for training to obtain good generalization performance by comparing the performance of models trained on various subsets of APLs (Figure 2f-g, Supplementary Figure 2b-c).

First, a given percentage of all APLs was randomly sampled as training data, while the remaining samples were used for testing (Figure 2f, Supplementary Figure 2c). In the educated repertoire, the average performance decreased only slightly when training on 25% of the available samples (*n* = 38 APLs) as compared to using 151 APLs for training: for regression, the Spearman correlation decreased by 0.041 at an effect size (mean difference over standard deviation) of 0.900, while for classification the AUC decreased by 0.026 at an effect size of 0.547. When further reducing the training samples to 10% (*n* = 15 APLs), a noticeable drop in performance was observed, resulting in a decrease in Spearman correlation of 0.147 (effect size: 2.976) and of 0.087 (effect size: 1.700) in AUC. The naïve repertoire followed a similar pattern, albeit with decreased initial performance. The performance remained stable until training on only 25% (*n* = 38 APLs), however, with a larger decrease in performance (Spearman correlation: 0.078, AUC: 0.051) compared to the educated repertoire. This was followed by a stronger decrease (Spearman correlation: 0.186, AUC:0.137) when training on fewer data (*n* = 15 APLs).

In order to gain insights into the interaction between epitope features and predictions, we further evaluated the model in two additional cross-validation settings by splitting the data either by amino acid or by position. In the first setting (leave-amino-acid-out, LAO), all APLs containing a given amino acid, in turn, were reserved for validation, while in the second setting (leave-position-out, LPO) the process was repeated based on epitope position instead of amino acids (Figure 2g, Supplementary Figure 2b). In the LAO setting, regression performance changed only negligibly for both educated (Spearman: 0.015, effective size: 0.291) and naïve repertoires (Spearman: −0.017, effective size: 0.085) suggesting that the model could successfully leverage the physiochemical features used to encode amino acids and, thereby, generalize to unseen amino acids with properties similar to those in the training set. However, when mutations at specific positions were not included in the training set, the model was barely able to predict T cell activation scores: the Spearman coefficient dropped to 0.057 across both repertoires, indicating hardly any correlation between predictions and observations. Similarly, the AUC decreased to 0.495, indicating random classification predictions. This highlights the importance of sampling across all epitope positions when predicting the mutational effects, and can be explained by the functional role of the different epitope positions [19]. While anchor positions fix the epitope within the MHC binding groove, and are therefore not accessible to the TCR, other positions are presented to the TCR to varying degrees and form the majority of interactions.

### Accurate prediction of the reactivity landscape for unseen TCRs

The previous experiments predicted T cell activation within an individual TCR, i.e., the model predicts the effect of mutations on a TCR for which several APLs were observed during training. As a next step, we further evaluated our model’s capability to generalize to new TCRs by predicting the effect of all APLs on an unseen TCR that was held out during training, following the aforementioned cross-validation scheme stratified by TCR (i.e., leave-TCR-out) on the whole human dataset. In addition to the wild-type epitope and APL sequence, we here provided the model with the sequence representation of the TCR CDR3*α* and CDR3*β* encoded by the Atchley factors described above [17]. The TCRs were padded to the same sequence length through Multiple Sequence Comparison by Log-Expectation (MUSCLE) [20], which aligns multiple biological sequences while still preserving homology. Overall, this provided the model with a residue-level representation of the CDR3 regions from which common sequence features can be learned to generalize to related TCRs.

During classification for unseen TCRs, the AUC for the educated repertoire was 0.905 on average, with a standard deviation of 0.041, and varied between 0.965 (TCR ED8) and 0.811 (TCR ED5) (Figure 3a-b, Supplementary Figure 3, Supplementary Table 1). The performance for the naïve repertoire was considerably more variable than for the educated repertoire, with an average AUC of 0.620 at a standard deviation of 0.286 during classification. The three receptors with the smallest amount of activated APLs, TCR G6, TCR F5, and TCR H5, had AUC values below 0.5 (respectively 0.426, 0.242, and 0.219), while the AUC for TCR OT-I was 0.959. This discrepancy between the predictive performance of different TCRs is not surprising: while receptors with high reactivity from the educated and naïve repertoire interact with the APLs derived from SIINFEKL in a similar manner and are thereby predictable, low-reactive TCRs are likely to recognize different cognate epitopes, and hence follow widely different interaction patterns [21]. Overall, the model’s classification indicated by the AUC shows a strong statistically significant Pearson correlation of 0.911 to the wild-type activation (*p*-value= 2.4 10*^−^*^10^, Supplementary Figure 4). Hence, we conclude that due to the composition of the dataset, our model is particularly suited for TCRs that show a high affinity towards the wild-type epitope.

**Figure 3.**
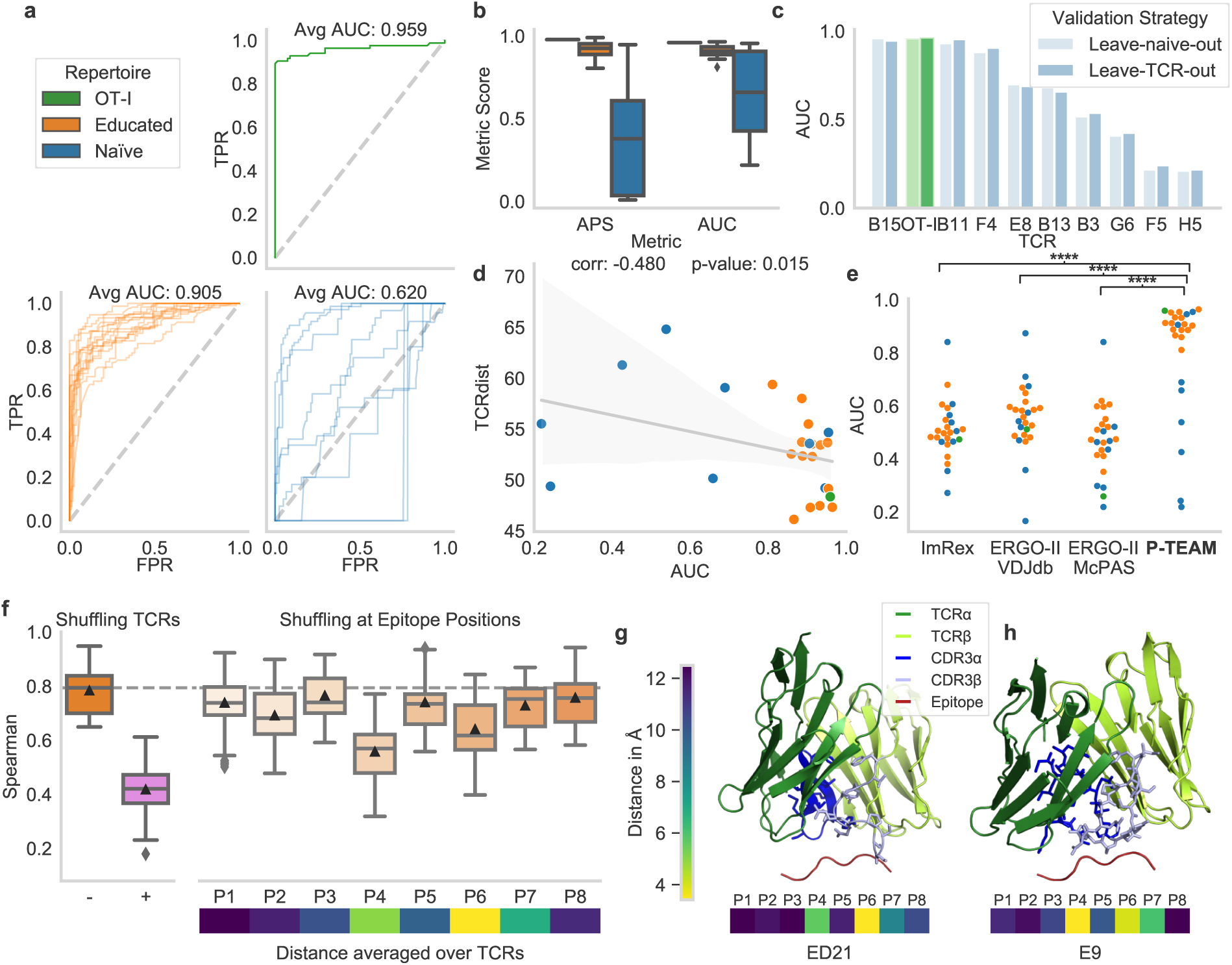
Generalization capabilities predicting activation for novel TCRs. **a**, ROC curves for the different groups of TCRs. **b**, Averages precision score (APS) and area under the ROC curve (AUC) as additional classification metrics. **c**, Comparison of the AUC scores when training on the full remaining murine dataset (Leave-TCR-out) or solely the educated repertoire (Leave-naïve-out). **d**, Classification performance shows negative correlation to the average TCRdist [22] between the training and test set. The boundaries of the linear regression line indicate the 95% confidence interval. **e**, P-TEAM significantly (****: p-value < 0.0001) outperforms existing TCR-epitope predictors ImRex [11] and ERGO-II [9]. **f**, the importance of input features obtained by replacing the test TCR input with a random CDR3 sequence of the dataset (+) or by shuffling the amino acid at each epitope position in the test set compared to the un-shuffled performance (- and dashed line). Below, the average distance of the center of mass between the epitope and TCR residues is shown. All box plots indicate the data quartiles while the whiskers extend to the extreme values excluding outliers. The median is indicated as a horizontal line and the mean as a triangle. **g, h**, Predicted structural model of the TCR and epitope, and minimal distance to the individual epitope positions for receptors ED21 and E9 (highest and lowest activation, respectively). The model shows the interaction between the epitope and the CDR3 of the TCRs.

To investigate the diversity in reaction patterns in the naïve repertoire, we trained the classification model on all TCRs in the educated repertoire and predicted the activation scores of the naïve repertoire. The resulting performance was compared with the earlier leave-TCR-out validation scheme. The classification performance measured by AUC scores obtained with these two methods were very similar with a maximal absolute difference of 0.025 (Figure 3c), providing additional evidence for our earlier conjecture that TCRs in the naïve repertoire were so diverse among each other that interaction patterns were not easily transferable from one TCR to the other.

To evaluate this further, we quantified the distance of each TCR towards the dataset by its mean TCRdist [22] to the remaining TCRs. The model’s classification performance for both repertoires indicated by the AUC showed a statistically significant negative Spearman correlation (*ρ* = 0.480, *p*-value=0.015, Figure 3d) to the TCR distance, indicating lower performance for less related TCRs. This analysis strengthens the evidence that P-TEAM performs well on reasonably similar TCRs, and its performance is reduced to the degree to which the TCRs are different. These results offer a reliable indicator to practitioners as to when our model is applicable and indicate the possibility of further improving the model’s performances by acquiring experimental data covering a broader spectrum of binding modes.

### P-TEAM outperforms conventional TCR-epitope predictors

Several machine learning approaches have been developed lately to directly classify pairs of TCRs and epitopes as binding or non-binding based on their sequences [6–11]. These tools are trained on curated databases of publicly available TCR-epitope pairs such as the IEDB [12], VDJdb [13], and McPas-TCR [23] databases, which, however, only contain a limited amount of epitope mutations. The different approaches largely vary in their machine learning techniques, the data used for training, and the input data the model receives about the TCR. Here, we report the comparative performance of the two common deep learning-based predictors ImRex [11] and ERGO-II [9], which provided trained models along with their publication. While ImRex utilizes a Convolutional Neural Network on a physiochemical sequence representation, ERGO-II relies on a pre-trained autoencoder composed of fully connected layers on one-hot-encoded sequences. We compared these predictors against the classification prediction of P-TEAM trained with the leave-TCR-out protocol (Figure 3e).

P-TEAM significantly outperforms ImRex, ERGO-II trained on VDJdb, and ERGO-II trained on McPas-TCR (paired two-sided *t*-test, all *p*-values<0.0001) by a large margin (increase in averaged AUC over all TCRs larger than 0.25). This confirms the previous observations that these models generalize poorly to unseen epitopes [24]. The mentioned databases used for training rarely contain data from deep mutational scans, which explains why the predictors did not learn to generalize to small changes in the epitope sequences causing large changes in activation patterns. Overall, most baseline models performed only slightly better than random on this challenging dataset (average AUC<0.60).

These results highlight that current general TCR-epitope classifiers cannot be used to predict T cell activation by APLs. This was expected as their training data did not contain many epitope mutations, and the model structure is not specifically designed to deal with small alternations in the epitope sequence. Until the number of diverse epitopes and mutational scans in public databases increases drastically, specialized datasets and predictors such as P-TEAM are therefore needed to study the effect epitope mutations have on T cell activation.

### P-TEAM learns biologically relevant interactions

To shed light on the inner workings of our model, we investigated the relevance of different input features in the most difficult predictive task we tested, i.e., predicting TCR reactivity scores for unseen TCRs in a regression setting. To perform a robust analysis, we employed permutation importance tests to assess feature importance, which is a standard technique in the toolbox of interpretable machine learning [25], and is especially suitable for our scenario, as our dataset contains all possible TCR-APL combinations [26]. After training the model on unperturbed data, the predictions were performed on datasets in which selected groups of input features were randomly shuffled, while the labels remained unperturbed. Intuitively, using random values for features that are important for prediction breaks their dependency on the target, and thus greatly reduces the model’s performance. Unimportant features, instead, are not used by well-performing models, thus using random values for them should not impact performance. We evaluated the model in two settings, using either a random, different TCR CDR3 sequence of the dataset, or permuting an epitope amino acid at a given position (Supplementary Figure 5). When replacing the full CDR3 region with a random sequence from the dataset during prediction, the performance decreased most noticeably by an absolute drop of 0.366 in Spearman coefficient in the educated repertoire. This behavior is expected and easily explained, as in this case, the model receives a different random TCR as input. Exemplary, assume the model was evaluated on TCR ED5 but the input was randomly perturbed to TCR ED8’s sequence. The activation patterns vary between both TCRs as TCR ED8 (in contrast to TCR ED5) is stable for mutations at epitope position P1 (Figure 2a). Hence, a generalized model cannot predict the effect of mutations on TCR ED5 based on TCR ED8’s sequence but must learn the reflective sequence features that are connected to the different activation patterns from different TCRs. As shown by this example, a large decrease in performance when shuffling the CDR3 sequences highlights that the model must have learned to incorporate the TCR information into its prediction and, therefore, showed the generalization ability of P-TEAM to adapt to novel TCRs. When analyzing epitope positions of SIINFEKL, the highest sensitivity was assigned to positions P4 and P6 with an absolute decrease in Spearman of 0.226 and 0.143 (Figure 3f). In contrast, epitope positions P1, P3, P5, and P8 remain robust to perturbations (decrease in Spearman < 0.05), indicating the low importance of the amino acid residues at these positions for TCR binding. In fact, the side chains at these positions of SIINFEKL have previously been reported as completely (P5, P8) or predominantly (P1, P3) buried within the groove of MHC class I H2k^b^ and, hence, are not in contact with the TCR [27]. The only exception is P2, which was also reported to be enclosed by the MHC but showed higher feature importance in our tests. Overall, it is biologically plausible that these positions do not play a significant role in TCR-epitope binding, and thus prediction thereof.

To further confirm that these results are in concordance with biological findings, we modeled the three-dimensional structure of each TCR and the epitope SIINFEKL using TCRpMHCmodels [28]. For each structure, the distance was calculated between the centers of mass for each amino acid residue of the epitope towards its nearest residue in the TCR. The sensitive positions P4 and P6 (Supplementary Figure 5a), which were previously identified protruding from the binding groove [29], laid in close proximity to the TCR with a distance less than 6 Ångström between the residues center of mass when averaging across the murine dataset (Figure 3f-h, Supplementary Figure 6), corresponding to a distance that indicates contact between residues [30]. Overall, the proximity of the epitope residues expresses a strong, significant correlation with the feature importance indicated by a decrease in Spearman correlation (Pearson: 0.722, p-value: 0.043, Supplementary Figure 7). The various TCRs of a dataset seem to engage with the pMHC in different binding modes (Supplementary Figure 6). Exemplary, the structural modeling of the TCRs ED21 (highest wild-type activation in the murine dataset), and E9, (lowest activation) revealed that both TCRs are in contact with positions P4 and P6 (Figure 3g-h). However, while TCR ED21 predominantly interacts with epitope position P6 via its CDR3*β* chain (as commonly detected in the educated repertoire, Supplementary Figure 6), TCR E9 lies in closer proximity to position P4, indicating additional involvement of the TCR *α*-chain.

Overall, the feature importance analysis of our model – combined with findings in the literature and our structural modeling – further validated the results obtained by P-TEAM. The model’s reliance on known biologically relevant features ensures that its good performance was not a statistical artifact but was based on learning the interaction between APLs and TCRs.

### Iterative experimental design decreases training set size to 24 APLs

So far, studying the effect of mutations entailed a comprehensive experimental evaluation of all possible APLs-TCRs combinations. As we showed above, however, P-TEAM could accurately predict T cell activation while being trained with as few as 25% randomly selected APLs (*n* = 38) in a classification and regression setting. In order to further reduce the wet-lab effort required to train P-TEAM on new TCR repertoires, we optimized the experimental design to find the smallest subset of APLs needed to learn a well-performing model, as opposed to the random selection of APLs tested above. Active learning [31, 32] is a collection of machine learning techniques that aim at iteratively improving a model’s performance, by deciding which samples to label experimentally (Algorithm 1). These techniques extend a small initial training dataset by requesting the label of one or more examples, whose knowledge is likely to improve the model’s performance the most. In this way, active learning can greatly reduce the total amount of data required to reach a desired performance, by only collecting diverse and informative examples for training. In practice, this procedure requires multiple experiments to be performed sequentially in the wet lab with APLs suggested by the P-TEAM active learning framework. However, the total amount of required samples, and therefore the total cost of collecting the dataset, is lowered through this alternating interplay between wet lab experiments and model training.

As our dataset already measured the effect of all possible APLs, we simulated this process by hiding the label for most APLs, and gradually, over *m* = 10 iterations, revealing the labels of a batch of examples (*n* = 8) whose prediction was most uncertain for the TCR-specific models (Methods Active learning). We compared our active learning method with a baseline that randomly chooses eight APLs to label in each iteration. To start the active learning procedure, we provided an initial training dataset consisting of one APL per position with the amino acid exchange that was most different from the wild-type epitope as quantified by the BLOSUM62 substitution probability [33]. Compared to a random selection, using this diverse initialization set (*n* = 8) improved the performance noticeably in the regression task by an increase in Spearman correlation of 0.120 in the educated repertoire (Figure 4a) reaching an average Spearman correlation of 0.614 from only eight observed samples. However, this is only reflected in a minor improvement during classification with an absolute increase of AUC of 0.024 (Figure 4b) at the first iteration. At iteration 10 (80 training APLS), the model achieves an AUC of 91.4%, which required a training set of 137 random APLs in previous experiments (Supplementary Figure 2c), thus reducing the amount of required training data by 42%. Overall, the active learning strategy statistically outperformed random sampling at every iteration on both datasets (unpaired *t*-test, *p*-value<0.0001 for regression and classification, with the exception of AUC at iteration 3, *p*-value=0.004).

**Figure 4.**
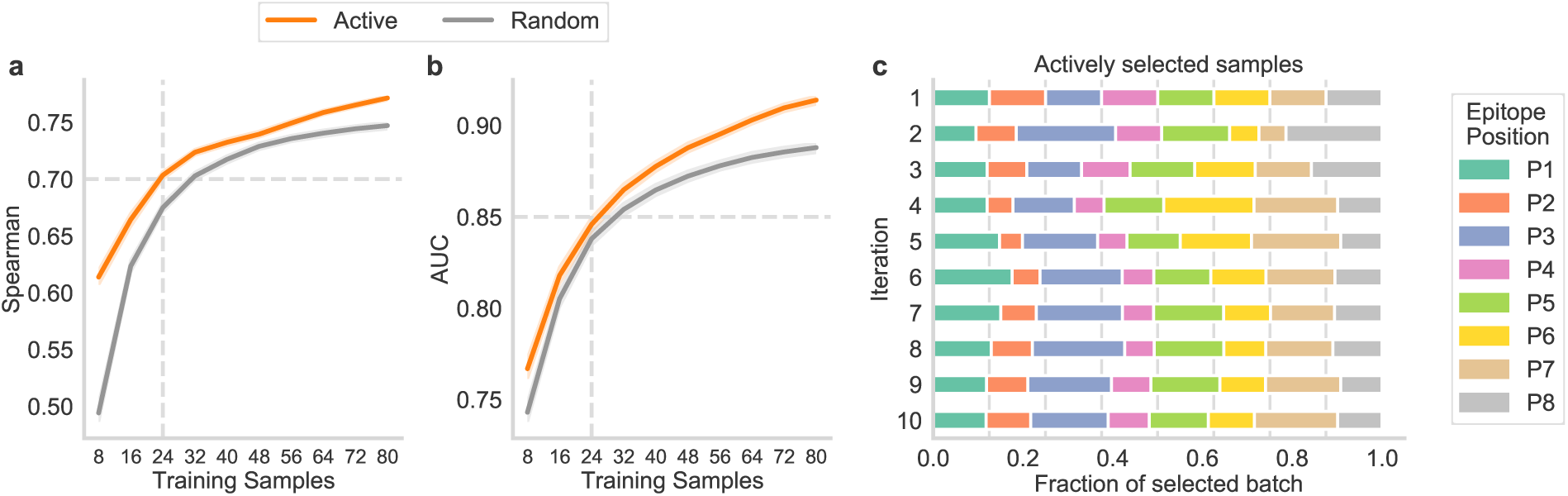
Reduction in training samples through active learning. Comparison of the active learning framework to random sample selection on the educated repertoire of the murine dataset for classification (**a**) and regression models (**b**) for predicting within a TCR. The expected performance is shown for up to *m* = 10 consecutive iterations (*n* = 80 APLs) of alternating wet lab experiments and model training. The dashed horizontal line indicates the performance threshold of 0.7 Spearman and 0.85, respectively, which can be obtained by using 3 iterations (24 APLs) of active learning as indicated by the dashed vertical line. The boundaries of the mean line indicate the 95% confidence interval. **c**, Fraction of the mutated positions of the APLs within the newly selected training batch during the active learning process for each iteration. The vertical lines represent a random selection of the samples.

This improvement over random sampling can be attributed to a dataset-specific focus on certain positions after a uniformly sampled initial set. P-TEAM expressed high uncertainty for mutations at positions P3 and P8 at the second iteration, leading to an oversampling with 22.1% and 21.4%, respectively, of the selected APLs stemming from this position compared to 12.5% for uniform sampling (Figure 4c). In the following iterations, samples were selected more uniformly. Over all 10 iterations, APLs with mutations at position P3 were sampled most frequently forming 17.1% of the training samples. Contrary, positions P2 and P4 were selected least with a frequency of 8.4% and 8.5%, respectively, as T cell reactivity was similar for all exchanges at these positions (Figure 2a,c). With this sophisticated interplay between experimental design and computational modeling, P-TEAM was able to reach a high performance of AUC greater 85% and Spearman greater 70% after the third iteration. We, therefore, conclude that three experimental rounds of alternating wet-lab and *in silico* experiments, collecting 24 APLs of an 8mer epitope (15.8% of the dataset) in total, are sufficient to train P-TEAM to a satisfactory performance level, which can be further improved with additional data collection.

### P-TEAM identifies cross-reactive APLs for neo-antigen specific human TCRs

So far, all experiments have been conducted on mutations of the murine model epitope SIINFEKL, for which the T cell response has been extensively characterized. To further validate P-TEAM on a therapeutically relevant epitope, we introduced a second mutational scan for the human cancer neo-antigen VPSVWRSSL with a similar structure as the murine dataset (Methods Data Collection, Supplementary Figure 8). The HLA-B*07:02-restricted epitope VPSVWRSSL occurs in a frameshift-induced neo-open reading frame (neoORF) of the gene RNF43 (unpublished data), which is frequently mutated in gastrointestinal cancers [34]. Seven TCRs with therapeutic potential were isolated from a naïve repertoire of VPSVWRSSL-binding T cells, of which six TCRs showed reactivity against the wild-type epitope and APLs. The human dataset contained the activation score between combinations of these six TCRs and 133 out of the 9×19=171 possible mutations (798 unique human pMHC-TCR interactions). The dataset did not contain APLs predicted to break MHC-binding via netMHCpan [35], which occurred especially for mutations at anchor positions P2 and P9 affecting 19 and 15 APLs, respectively. Contrary to the murine dataset, all human TCRs recognized at least 20 mutations with a maximum of 75 mutations (Figure 5a). However, the two TCRs R24 and R28 were less activated (Figure 5b) caused by limited reactivity at the end positions P7 to P9, while the activation score of the remaining TCRs is robust to exchanges at these positions (Figure 5a). Overall, the change in activation score was again highly dependent on the position of the mutation. Exchanges at center positions P4, P5, and P6 generally led to a drop in activation (Figure 5c).

**Figure 5.**
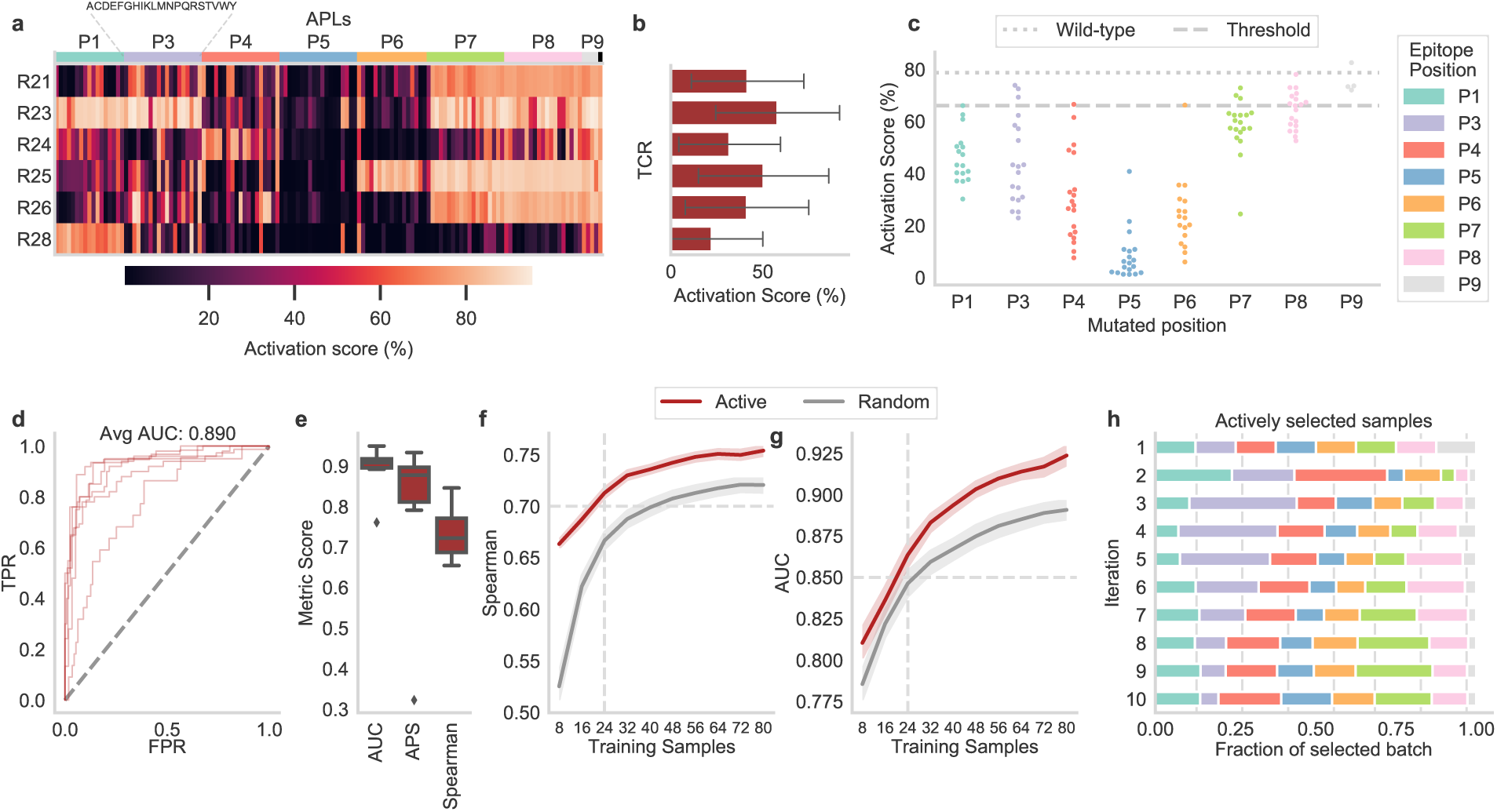
Predicting the effect of neo-antigen mutations within a human TCR. **a**, The normalized activation scores of six TCRs of the human dataset express high activation against the mutation landscape. **b**, The activation scores averaged for all APLs of one TCR indicate two reactivity patterns. **c**, The epitope position on which the mutation occurs strongly influences the activation per APL (*n* = 19, except P1: *n* = 17, P6: *n* = 17, and P9: *n* = 4) averaged over all TCRs (*n* = 6). The threshold value represents the boundary between binding and non-binding and wild-type indicates the activation scores of the base epitope VPSVWRSSL. **d**, The Receiver operating characteristic (ROC) curves the six human TCRs indicate the True Positive Rate (TPR) against the False Positive Rate (FPR) at all prediction values as thresholds. **e**, Different evaluation metrics for regression (Spearman) and classification models (APS: Average Precision Score, AUC: Area Under the ROC Curve). Comparison of the active learning framework to random sample selection on the human dataset for classification (**f**) and regression models (**g**). The expected performance is shown for up to *m* = 10 consecutive iterations (*n* = 80 APLs) of alternating wet lab experiments and model training. The dashed horizontal line indicates the performance threshold of 0.7 Spearman and 0.85, respectively, which can be obtained by using 3 iterations (24 APLs) of active learning as indicated by the dashed vertical line. The boundaries of the mean line indicate the 95% confidence interval. **h**, Fraction of the mutation positions of the APLs within the newly selected training batch during the active learning process for each iteration. The vertical lines represent a random selection of the samples.

We observed a similar performance to the educated murine repertoire when training P-TEAM for individual human TCRs. During LMO classification, P-TEAM achieved an average AUC of 89.0% and APS of 78.4% (Figure 5d-e, Table 2) with the worst performing TCR R24 (AUC: 76.1%, APS: 32.2%) and the best performing TCR R25 (AUC: 94.9%, APS: 93.3%). Predicting the activation score in the regression setting also expressed a high performance with a Spearman correlation of 73.4% (Figure 5e) in the human dataset compared to 76.0% in the educated murine repertoire (Figure 2e) indicating that the approach can be applied to different epitopes as well as human TCRs.

As in the murine dataset, the model performed only slightly worse when trained on 25% of the human APLs (*n* = 33, Supplementary Figure 9a-b) with a decrease of 0.044 in Spearman correlation (effect size of 0.605) and a decrease of 0.024 in AUC (effect size 0.305), followed by a large drop in performance when training on fewer data (Spearman correlation: 0.135 at effect size 1.527, AUC: 0.073 at effect size 0.833). Again, active learning further improved sample efficiency for the human dataset (Figure 5f-g). With the initial set of only *n* = 8 samples, the model reached a Spearman correlation of 0.663 and an AUC of 0.810, outperforming random sampling by 0.138 and 0.025, respectively. Following, the model steadily improves to a Spearman correlation of 0.754 and an AUC of 0.924 at iteration 10 (*n* = 80 APLs), significantly outperforming random sample selection at every iteration (unpaired *t*-test, *p*-value<0.0001 for classification and regression, with exception of AUC at iteration 1, *p*-value=0.015). While in the murine dataset the difference diminished in the regression setting for later iterations, active learning surpasses random sampling at all steps by a margin of at least 0.03 in Spearman correlation except for iteration 8 (0.029). At the threshold of *n* = 24 APLs determined in the murine dataset, P-TEAM was here able to predict T cell activation with an AUC of 0.864 and a Spearman correlation of 0.713. As in the murine dataset (Figure 4c), APLs with exchanges at Position P3 were identified as beneficial to the training process with a frequency of 19.5% at the first iteration and 18.1% in total (Figure 5h). However, positions P1 and P4, which were seldomly selected in the murine dataset at iteration 1 (9.5% and 10.3%, respectively), were heavily oversampled with 23.9% and 28.9% in the human dataset. This focus on partially different positions between both datasets further highlights the necessity that an uncertainty-based experimental design is required as no generalized rules can be derived across different epitopes.

While the performance on the human dataset in most previous experiments generally followed the trend of the educated murine repertoire, the average model’s performance was reduced (AUC: 0.771, APS: 0.663, Spearman: 0.663) for predictions in the Leave-TCR-out setting (Figure 6a-c). Specifically, the model failed to generalize to the two TCRs R24 and R28 with an AUC of 0.561 and 0.618. Both TCRs were experimentally identified to follow different activation patterns at positions P7 to P9 (Figure 5a). Presumably, a larger variety in high-affinity binding modes needs to be captured within the repertoire data in order to further improve the capability to generalize to unseen TCRs with varying activation patterns. Despite the low performance on these two TCRs, the model out-performed the general TCR-epitope predictors ImRex and ERGO-II by an increase in AUC of 0.116 to 0.289 (statistically for ERGO-II VDJdb at a *p*-value: 0.014, and ERGO-II McPas *p*-value: 0.005, at *n* = 6, Figure 6d).

**Figure 6.**
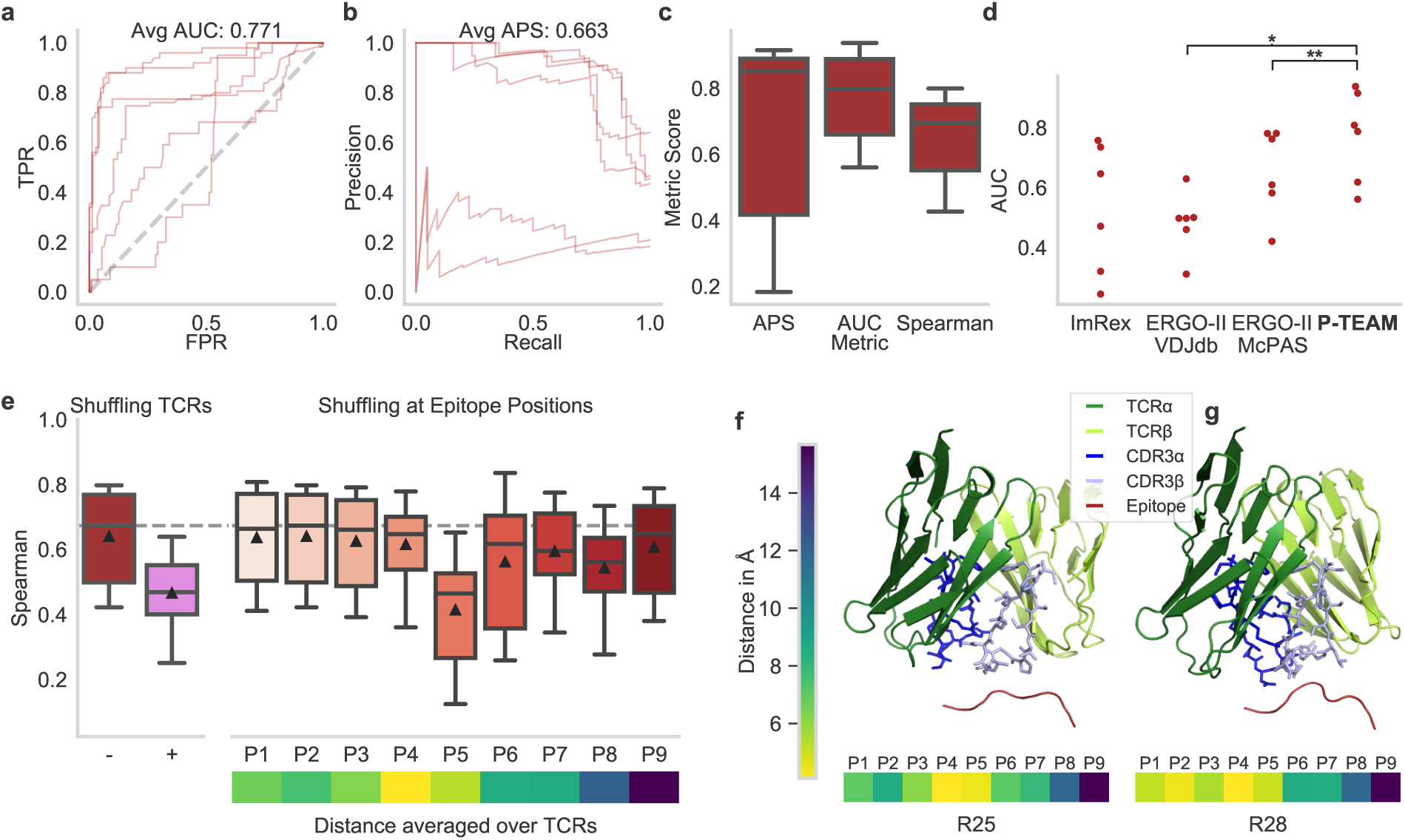
Across-repertoire prediction for the human dataset. ROC (**a**) and Precision-Recall (**b**) curves for the six human TCRs. **c**, Averages precision score (APS), area under the ROC curve (AUC), and Spearman correlation as classification and regression metrics. **d**, P-TEAM outperforms existing TCR-epitope predictors ImRex [11] and ERGO-II [9] by a large margin (*: p-value < 0.05, **: p-value < 0.01). **h**, the importance of input features obtained by replacing the test TCR input with a random CDR3 sequence of the dataset (+) or by shuffling the amino acid at each epitope position in the test set compared to the un-shuffled performance (- and dashed line). Below, the average distance of the center of mass between the epitope and TCR residues is shown. All box plots indicate the data quartiles while the whiskers extend to the extreme values excluding outliers. The median is indicated as a horizontal line and the mean as a triangle. **e, g**, Predicted structural model of the TCR and epitope, and minimal distance to the individual epitope positions for receptors R25 and R28 (highest and lowest activation, respectively). The model shows the interaction between the epitope and the CDR3 of the TCRs.

In contrast to the murine dataset, feature importance is here predominantly assigned to epitope position P5 (decrease in Spearman: 0.226, Figure 6e) that lay with an average distance of 5.64 Ångström second closest to the TCR in the structural models obtained by TCRpMHCmodels [28] (Figure 6f-g, Supplementary Figure 10). The effects of the anchor positions could not be observed as the dataset contained only four of the 19 possible APLs with mutations at P9 and none for P2 since the remaining mutations at these positions were predicted to break MHC-binding and were therefore not experimentally determined. However, P3, which was ranked second last in importance (decrease in Spearman: 0.024), had been reported to form an optional stabilizing interaction to the MHC class I HLA-B07*02 [36]. The predicted spatial models show that the two outlier TCRs R24 and R28 also follow different structural patterns. Both TCRs lay closer to positions P1 and P2 than the remaining TCRs (Figure 6f-g, Supplementary Figure 10), which might further indicate a different TCR-pMHC interaction pattern. However, it must be noted here that structural interpretation on the human dataset must be viewed with caution as the underlying template epitope in TCRpMHCmodels [28] expressed only 33.3% sequence identity to VPSVWRSSL.

To summarize, P-TEAM achieved high performance in predicting the effects of mutations in the neo-antigen VPSVWRSSL on T cell reactivity during classification and regression, even though the Leave-TCR-out setting is slightly limited due to the low amount of available TCRs. The model adapted to the changing effects of mutations at specific epitope positions as shown through changes in feature importance and differing sample selection during active learning. Based on the results of this novel dataset, we, therefore, conclude that P-TEAM can be used for biologically and therapeutically relevant TCRs across different host organisms, epitopes, and MHC alleles if a sufficient amount of annotated samples is available.

## Discussion

Antigen recognition by T cells is a fundamental step to enable the adaptive immune response. For this reason, pathogens and cancer cells try to escape surveillance by the immune system through epitope mutations that prevent TCR binding. Indeed, single point mutations can be enough to evade previously formed immune memory [37]. Understanding and predicting the effect of such mutations on TCR binding and T cell activation is thus essential to develop the next level of vaccines and immunotherapies. TCR binding prediction, however, is still an open problem due to the enormous variety of receptors and epitopes as well as the scarcity of paired datasets. Predicting the effect of point mutations is especially hard, as public datasets contain very few examples of epitopes differing by one residue towards the same receptor, making methods trained on these datasets not suitable for predicting the effect of point mutations.

As a first step towards this goal, we here introduce P-TEAM, a single-point mutational effect predictor trained on two novel datasets that measured TCR reactivity levels for single-point mutations of the wild-type model epitope SIINFEKL [15]. Importantly, the data were contextualized by assigning a defined reactivity value that leads to effective *in vivo* recruitment of T cells into an immune response [15]. We modeled the interaction of T cells to epitope mutation for individual TCRs – as well as across repertoires – with high accuracy, indicating the validity of our approach for different epitopes and host organisms. The model was able to learn this interaction based on the APL and TCR CDR3 sequences even when trained on a limited amount of annotated samples. While most prediction methods treat the TCR-epitope interaction as a binary event of recognition or non-recognition, P-TEAM could not only predict such a classification but also a continuous reactivity score in a regression setting.

We found that the model’s predictions were driven by a few highly sensitive residues in the epitope, and were not affected by changes in others. The model’s focus on specific APL positions was compared to structural TCR-epitope models to confirm its biological relevance. Based on the predicted spatial proximity of epitope and TCR residues, we validated that the model extracted meaningful interactions of the TCRs to residues of the APLs, which further are in line with previous findings in the literature. TCR-specific models could be trained with one quarter of all possible mutations (*n* = 38) without any notable changes in prediction performance. We further proposed an iterative experimental design framework that optimizes the collection of training samples for novel TCRs by an alternating process of wet lab experiments and model training. With this active learning framework, only 24 samples (15% of the APLs) were required to train P-TEAM in order to generate good predictions.

Finally, we applied P-TEAM to a dataset of therapeutically-relevant TCRs reactive to the human cancer-derived neo-epitope VPSVWRSSL (newly collected data). While observing similarly high performance as in the murine dataset, we observed epitope-specific differences in feature importance and sample selection, which highlight the adaptability of P-TEAM to novel epitopes and host organisms. Overall, P-TEAM makes it possible to significantly reduce the cost and effort of wet lab experiments to identify mutations that result in cross-reactivity, thus making applications such as the development of T cell-based immunotherapies cheaper and faster to develop.

Our modeling approach can be extended in several ways, for example by using the V(D)J-gene type as model input to guide the predictions. Together with more and more data, this will ultimately improve generalization across TCRs that are less related. Generalization could also be improved by using pretrained TCR embeddings obtained from models such as TCRbert [38]: through representation learning, we could harness larger databases of unpaired receptors to learn more accurate TCR similarity functions and thus guide P-TEAM towards better-generalized predictions when predicting activation for unseen TCRs. Finally, recent advances in structural modeling sparked by AlphaFold2 [39] and similar methods could be harnessed to create holistic binding models analyzing the TCR-peptide-MHC complex interactions.

In conclusion, we present P-TEAM, a TCR-epitope binding predictor specialized in single-point mutations of epitopes that generalizes across receptors and can be trained with as few as 24 mutations. The model is able to estimate continuous activation which ultimately characterizes TCR-epitope interactions beyond binary recognition. Our findings point to the intriguing possibility of predicting changes in T cell functionality due to single-point mutations in a quantitative manner from epitope and TCR sequence alone. P-TEAM thereby bears the potential of improving the safety and effectiveness of immunotherapies and vaccines.

## Code availability

The code including all experiments to reproduce the results, analysis, and tutorials is available at https://github.com/SchubertLab/TcrPrediction_MutatedAPLs.

## Data availability

All data including the murine and human datasets, aligned TCR sequences, and structural models used in this paper were made publicly available and are included in the Supplementary Material.

## Author Contributions

B.S. and K.S. conceived the project. E.D. and F.D. performed research, implemented models, and performed analysis. A.S., P.H., K.I.W, and D.H.B. acquired the experimental data. B.S., K.S., D.H.B., and B.B. supervised the research. All authors wrote the manuscript.

## Acknowledgments

E.D. and F.D. are supported by the Helmholtz Association under the joint research school “Munich School for Data Science - MUDS”. F.D. acknowledges financial support from the Joachim Herz Stiftung. K.S. is supported by the BMBF (project 01KI2013). P.H. is supported by the Else Kröner-Stiftung (project 2020_EKEA.127) This work was mainly supported by the BMBF grant DeepTCR (project 031L0290A) to K.S. and B.S.

## Methods

### Data Collection

#### Murine dataset

In this work, we analyzed the dataset described in Straub et al [15]. The authors experimentally determined TCR functional reactivity in response to mutations of the SIINFEKL epitope presented on the H2K^b^ allele. Each epitope residue at every position was exchanged against all other 19 encoded amino acids, at a time, resulting in a library of 152 unique mutations of the wild-type peptide. Functional reactivity against these APLs was experimentally determined for 36 different murine TCRs as described by Straub et al [15]. In brief, Jurkat triple parameter reporter cells (JTPRs) were engineered to express a single SIINFEKL-reactive TCR, and co-incubated with peptide-pulsed splenocytes. After 20h incubation time, NFAT reporter expression was assessed via flow cytometry. The murine TCR library consisted of 15 unique TCRs isolated from the memory compartment of mCMV-SIINFEKL infected C57Bl/6 mice (educated repertoire, Supplementary Data 1) [16, 40], as well as 20 SIINFEKL-reactive TCRs from a naïve C57Bl/6 donor (naïve repertoire, Supplementary Data 2) [15]. The TCR OT-I was included as a reference control. TCR sequences were isolated from single-cell sorted, H2k^b^/SIINFEKL multimer positive CD8^+^ T-cell clones stemming from either infected or naïve donors via the TCR SCAN platform [41] as described by Straub et al. TCR functional reactivity as assessed by JTPR stimulation was normalized as an activation score *A* across experiments (Supplementary Data 1-2). JTPRs expressing a unique TCR were stimulated with the APL library in independent experiments resulting in NFAT*_apl_*. In order to normalize the data, JTPRs of each TCR were included simultaneously in a single experiment and stimulated with the wild-type peptide resulting in NFAT*_sim_*. NFAT*_AP L_* expression from APL library stimulated JTPR were normalized to this experiment:

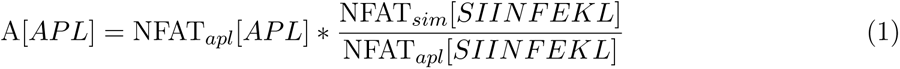

For computational analysis to predict meaningful T cell activation, we set a threshold in the activation score of 46.9%. As described by Straub et al., this value was experimentally determined in this screening platform to predict effective recruitment and clonal expansion *in vivo* after adoptive transfer of low numbers of TCR transgenic naïve T cells and infection. For the predictions, we excluded five TCRs for which the activation scores of all APLs fall short of this threshold.

#### Human dataset

This dataset was experimentally generated in an analogous manner to the murine dataset. The APLs were formed by every single-amino acid mutation of the human cancer neoepitope VPSVWRSSL. Prior to the experiments, binding of the APLs to the HLA-B*07:02 allele was computationally determined via NetMHCPan 4.1 [35]. APLs without predicted binding were excluded from the dataset affecting positions 1, 2, 6, and 9 with 2, 19, 2, and 15 mutations, respectively, leading to a total amount of 133 peptides. JTPRs were engineered to express a single human TCR recognizing the VPSVWRSSL epitope. JTPRs were co-incubated with peptide-pulsed K562 cells expressing HLA-B*07:02 for 20h. After 20h incubation time, NFAT reporter expression was assessed via flow cytometry. The repertoire consists of 7 TCRs isolated via pMHC multimer staining and antibody staining for a naïve phenotype (CD3^+^ CD8^+^ CD45RA^+^ CD62L^+^) from healthy donors. In all experiments, the TCR R27 was excluded as it did not show the expected activation against all APLs. The percentage of activated cells was experimentally determined for the remaining combinations leading to a total of 798 pMHC-TCR interactions. The activation scores were normalized by their positive control. 66.09% was chosen as a threshold for binarization, which represents the lowest activation of a TCR against the cognate epitope alongside sub-optimal tumor-cell lysis *in vitro* (unpublished data, Supplementary Data 3).

### Predictors

#### Data Representation

To provide the Random Forests with information on the APLs, we encoded their sequences into numeric representations. To this end, we represent each amino acid via five factors representing a summary of physiochemical properties as developed by Atchley et al [17]. Based on this encoding, we provided the full APL sequence and the difference between the APL and the wild-type sequence. Additionally, the position of the mutation, the original, and the new amino acid at this position are provided.

When predicting across TCRs, we provided the CDR3 sequences of the *α*- and *β*-chain as additional input to the Random Forests. To this end, we represented the amino acid of each position via the Atchley factors as described above. To counteract the effect of different lengths, we aligned the sequences using the MUSCLE algorithm [42] for the murine and human datasets, separately (Supplementary Data 4). Padding tokens were consequently encoded with all zero values.

#### Random Forests

The Random Forest predictors were consequently trained on this representation. While each Random Forest consisted of 250 different Decision Trees when predicting activation for the remaining APLs of a TCR, the number of Decision Trees was increased to 1,000 for cross-TCR prediction to tackle the larger feat*_√_*u<u>r</u>e space. Each individual tree was fit on a bootstrap sample of the data and a random subset of *d* features. We chose this setup with a large amount of diverse trees to prevent overfitting and aid generalization [43], and the resulting performance is saturated by the number of trees (Supplementary Figure 11). The trees were fully grown using the Gini impurity as the splitting criterion in case of classification and mean absolute error for regression, in order to avoid outliers dominating the models’ predictions.

### Baseline TCR predictors and distances

The data for the TCR predictors ERGO-II [9] and ImRex [11] were formatted as described by the authors in the corresponding GitHub repositories. While the trained model provided for ImRex uses the CDR*β* sequence as a sole input, ERGO-II can optionally incorporate the CDR3*α* sequence, V- and J-genes of both chains, and MHC type. To allow fair comparison, we reduced the input for ERGO-II to the information used by P-TEAM, i.e. the sequences of the CDR3 sequences of both chains. ERGO-II offers two different models which were trained on the VDJdb [13] and McPas-TCR [23] databases, respectively. Since it is unclear which model better fits the data used in this work, we report the performance of both models. Distances between the TCRs within a dataset were calculated based on the implementation of TCRdist3 [22] (Supplementary Data 5).

### Structural Modelling

The full nucleotide sequences of the TCR*α* and TCR*β* chains (Supplementary Data 1-3) were translated to amino acid sequences. *α*- and *β*-chains of each TCR, the wild-type epitope and the sequence of the MHC (H2k^b^ for SIINFEKL, HLA-B*07:02 for VPSVWRSSL) were used as input for TCRpMHCmodels-1.0 [44] to derive the structural models (Supplementary Data 6-7). Four TCRs of the educated repertoire (TCR ED5, TCR ED10, TCR ED23, and TCR ED40) were excluded from the following tasks, as the modeling software failed to derive structural models presumably due to lack of TCR templates for these sequences. These models were aligned by their MHC and visualized with PyMol [45], which also served as an interface to determine the structural relationships. The distances of the center of mass for each amino acid residue in the CDR3*α* and CDR3*β* towards all peptide residues were calculated via the ‘centerofmass’ command. Following, the minimal distances between TCR and epitope were determined for each epitope position.

**Algorithm 1.**
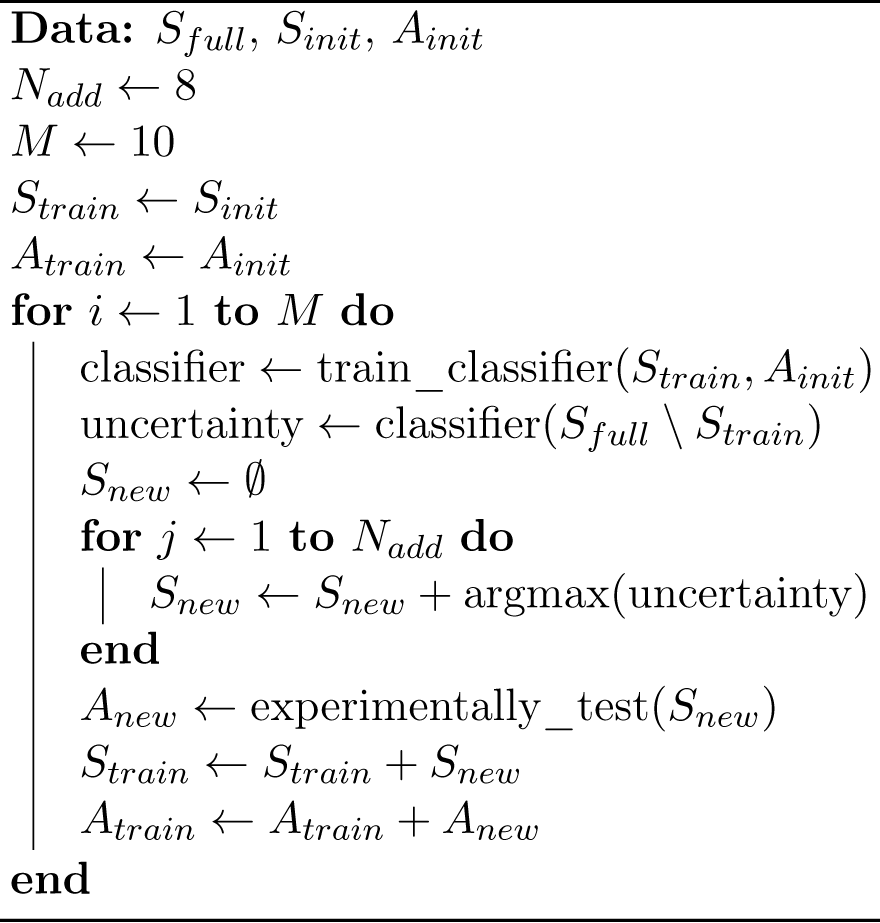
Sample selection of P-TEAM.

### Active learning

Active learning was used to reduce the amount of data needed to derive accurate predictors by choosing training samples in a sophisticated manner. We applied active learning in two settings. First, the algorithm selected the best APLs for an individual TCR to predict the remaining APLs. Second, given a set of TCRs for which the activation score was known for all APLs, the algorithm selected the best APLs to be experimentally determined for a novel TCR. The general workflow of the active learning procedure followed an iterative approach (Algorithm 1). In our experiments, we simulated the iterative experimental procedure by holding out the activation scores for all yet unknown samples.

#### Initialisation

The activation scores *A_init_* were experimentally determined for an initial set of training samples *S_init_* out of the full set of samples *S_full_*. This initial set consisted of the APLs with the largest BLOSUM62 [33] distance to the base epitope for each position and the wild-type epitope itself. These *S_init_* and *A_init_* were assigned as training samples *S_train_* and training activation labels *A_train_*.

#### Iterative process

After initialization, the iterative process is started. A classification predictor following the same model as described above was trained on *S_train_* and *A_train_*. This classifier predicted the binary activation for the remaining APLs (*S_full_ S_train_*). In each step, the *N_add_* APLs *S_new_* with the most uncertain prediction were identified and the corresponding activation scores *A_new_* were experimentally determined. Following, *S_new_* and *A_new_* were added to *S_train_* and *A_train_*, respectively. After the evaluation of the yet unobserved data, the iterative process continued with this updated training set until *M* = 10 iterations were reached.

#### Uncertainty

This active learning process requires a measure of prediction uncertainty for each sample. Since the Random Forest consisted of an ensemble of different Decision Tree classifiers, the proportion of votes between these individual predictors can be interpreted as the class probability of the Random Forest. Since the models were biased towards the dominating class of the training set, the inverse difference of this class probability for each sample towards the average class probability across all samples was used to indicate the uncertainty of the model.

#### Evaluation

At each iteration, the predictors were tested for classification as well as regression based on the selected samples *S_train_*. For this evaluation, the unobserved APLs (*S_full_ S_train_*) were used. The active learning scheme was compared against a baseline model, for which training data was added randomly. The experiments were conducted on 100 random seeds to obtain robust performance estimates for the different acquisition methods.

## Supplementary Figures

**Supplementary Figure 1.**
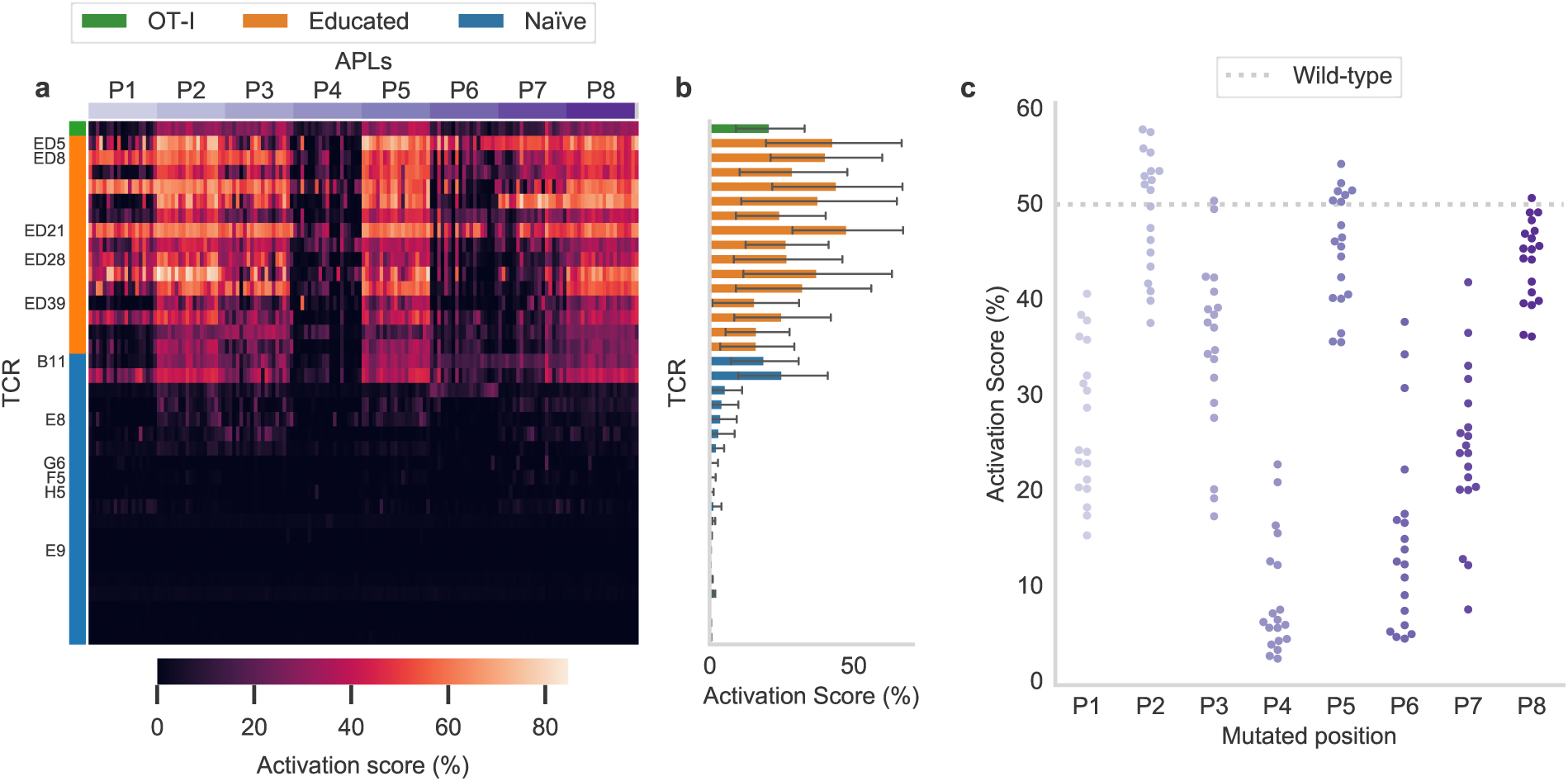
Unnormalized T cell activation for the TCRs and APLs of the murine dataset. **a**, Unnormalized activation scores. **b**, Unnormalized activation scores averaged for all APLs per TCR. **c**, Unnormalized activation per APL over all TCRs.

**Supplementary Figure 2.**
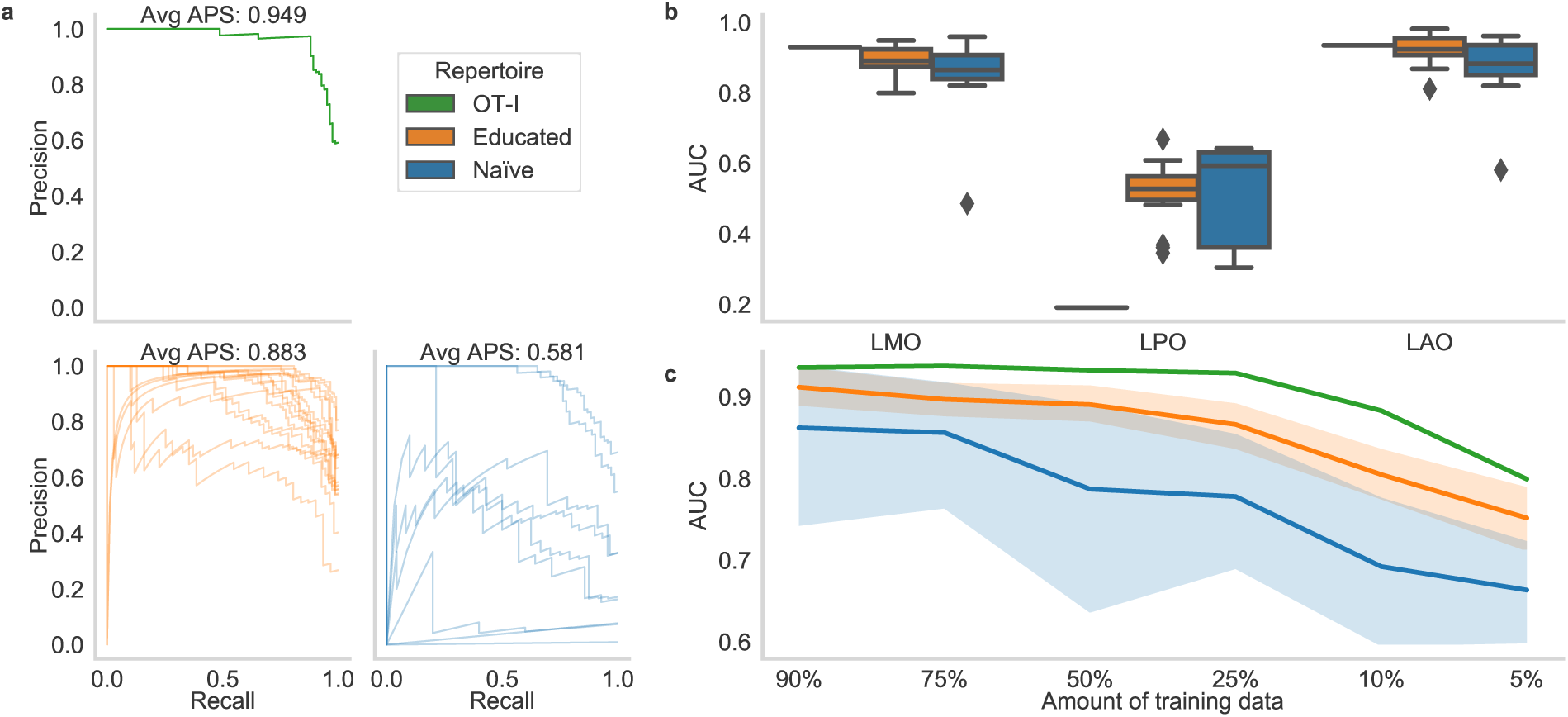
Additional metrics for predicting within a TCR of the murine dataset. **a**, Precision-Recall curve with Average Precision Scores (APSs). **b**, AUC obtained when training on different subsets of the data. LMO: leave-mutation-out, LPO: leave-position-out, and LAO: leave-amino-acid-out. The box plot indicates the data quartiles while the whiskers extend to the extreme values excluding outliers. The median is indicated as a horizontal line. **c**, AUC when a smaller amount of training data is used (average over ten repetitions with random subsets for each TCR). The boundaries of the mean line indicate the 95% confidence interval.

**Supplementary Figure 3.**
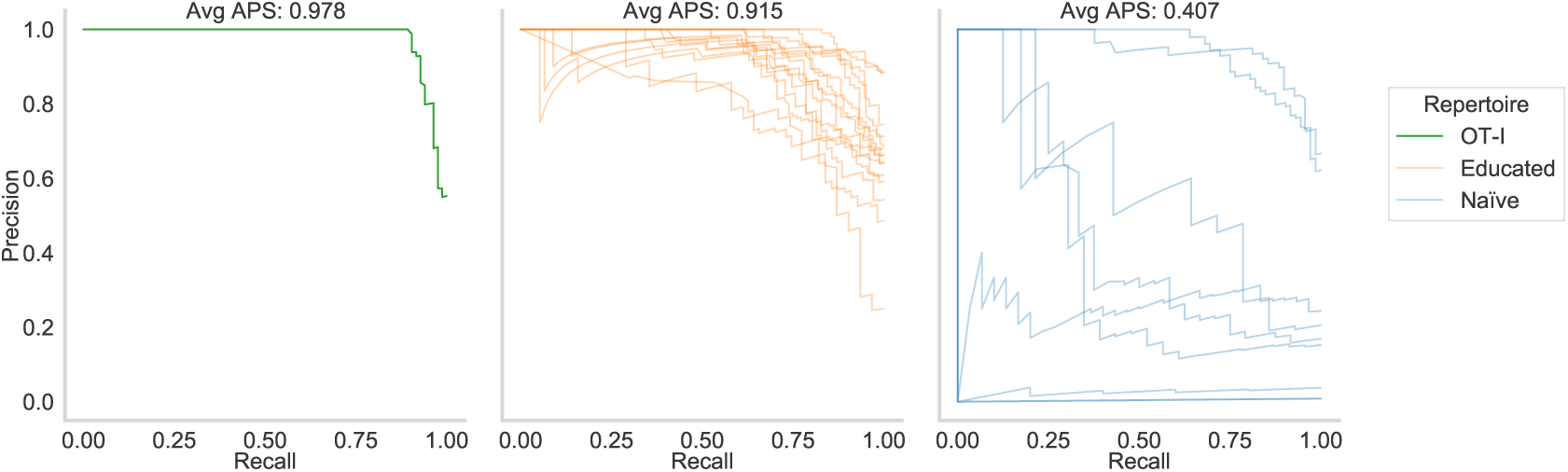
Additional metrics for predicting across TCRs of the murine dataset. Precision-Recall curve with Average Precision Score (APS) for the different groups of TCRs.

**Supplementary Figure 4.**
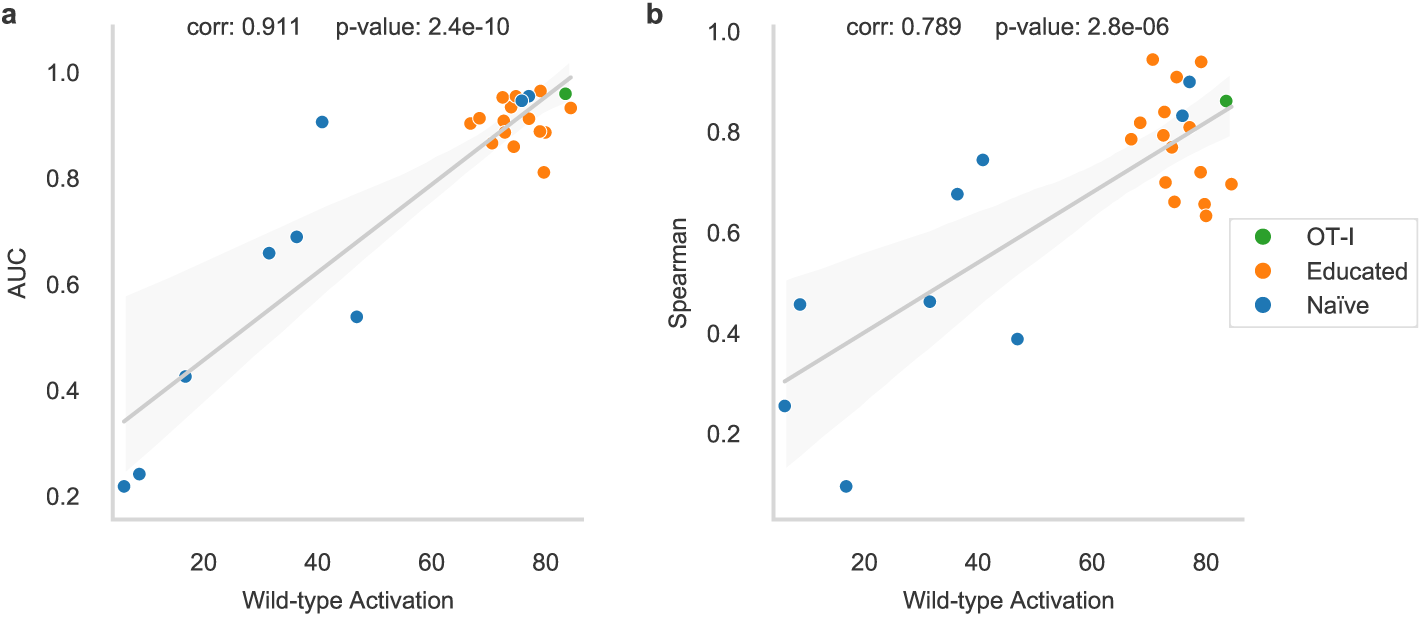
Relationship between generalisation performance and wild-type activation. The performance for classification (**a**) and regression (**b**) in the Leave-TCR-out setting shows a strong Pearson correlation to the normalized activation score against the wild-type epitope in the murine dataset. The boundaries of the linear regression lines indicate the 95% confidence interval

**Supplementary Figure 5.**
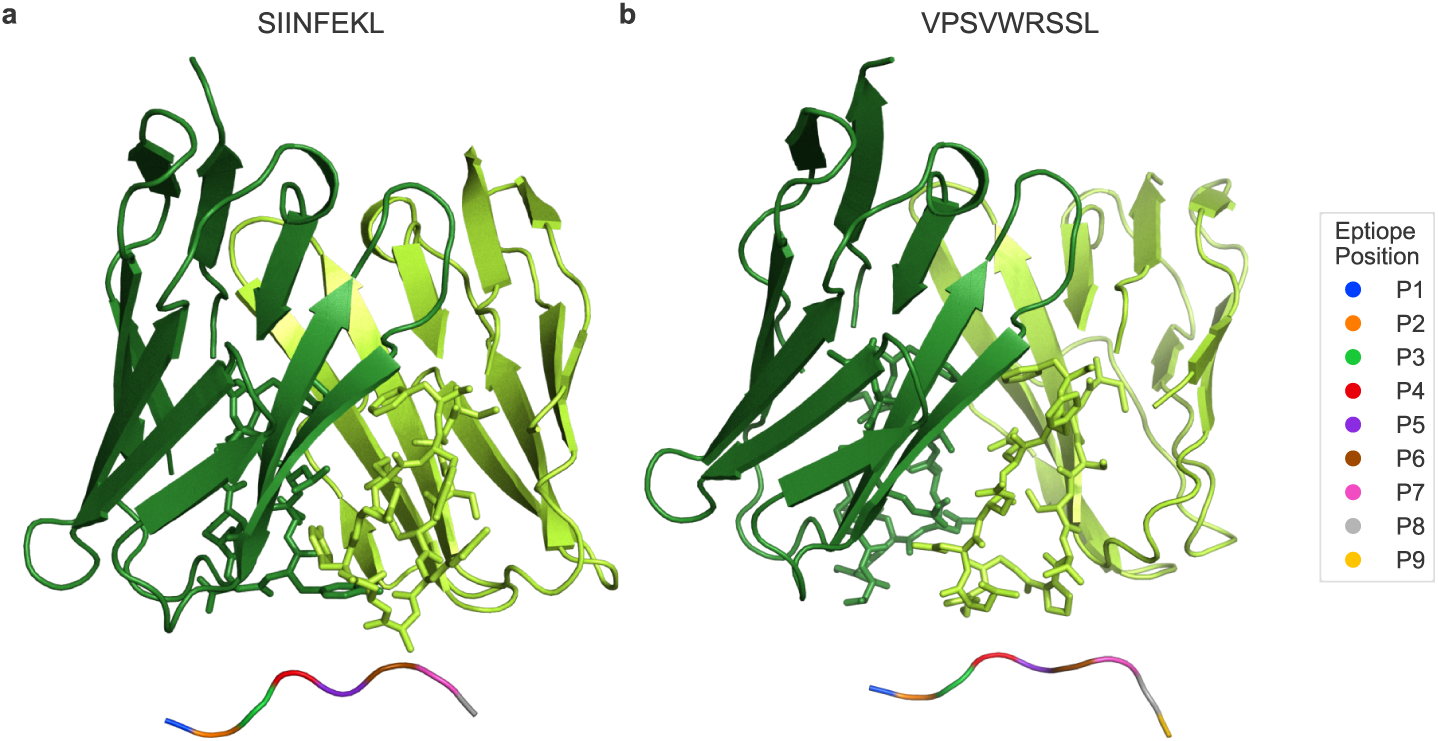
Epitope positions in the structural models. Epitope positions are highlighted in the structural models for OT-I/SIINFEKL (**a**) and R25/VPSVWRSSL (**b**). TCR*α* and TCR*β* chains are shown in dark and light green, respectively.

**Supplementary Figure 6.**
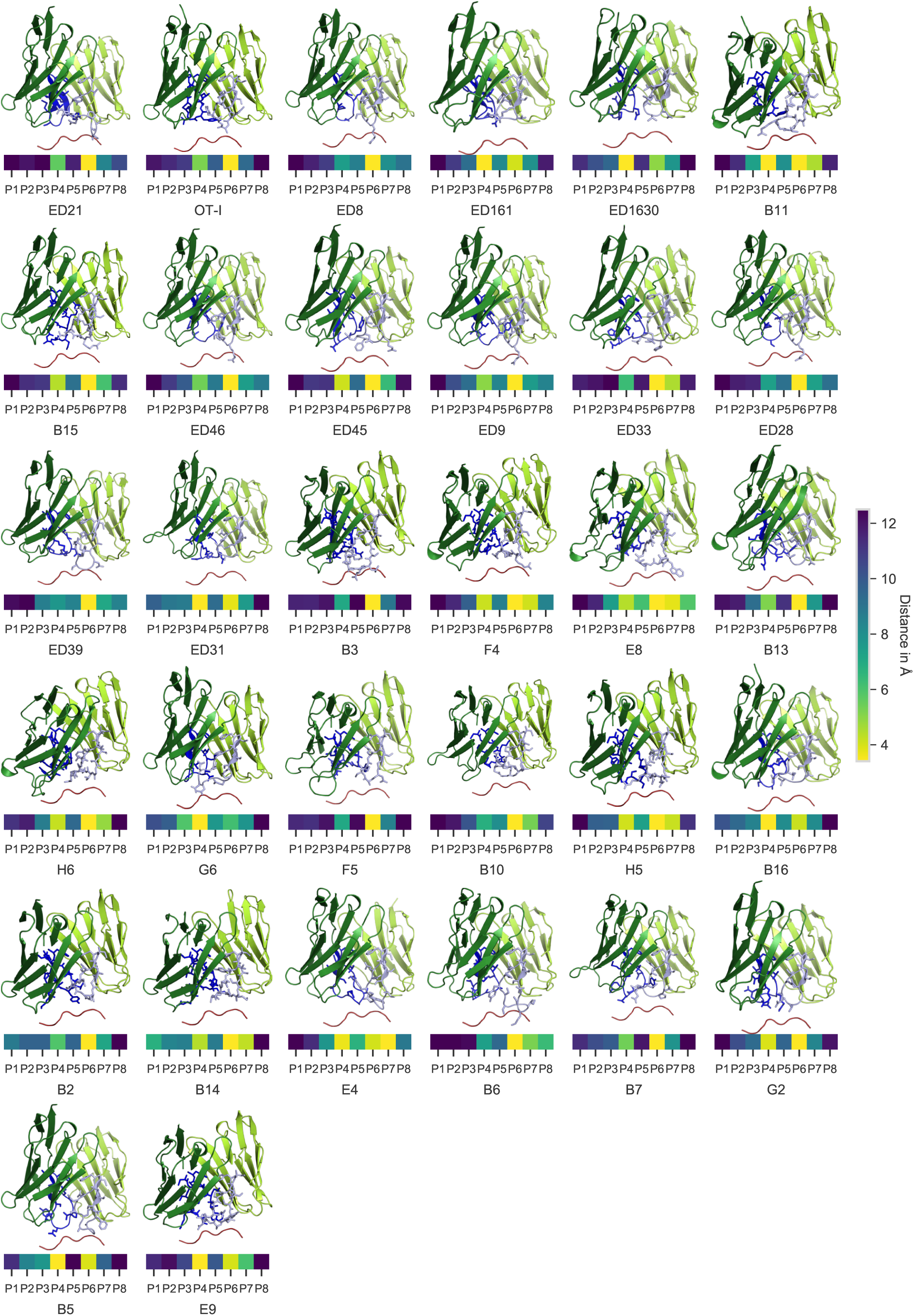
Structural Models of the murine dataset. Predicted structures of the TCR and epitope, and minimal distance to the individual epitope positions for all receptors of the murine datasets ordered by descending activation to the wildtype epitope.

**Supplementary Figure 7.**
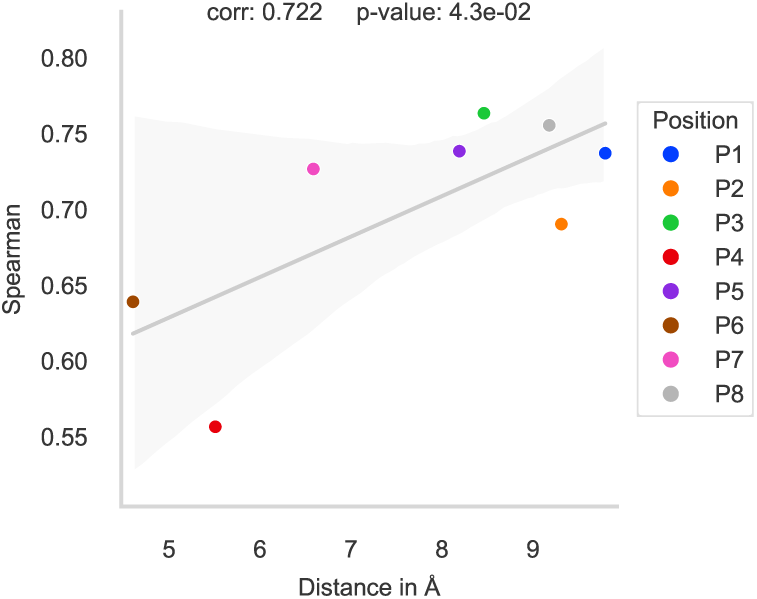
Relationship between spacial distance and feature importance. The regression performance during perturbation tests at each epitope position shows strong Pearson correlation to the distance between this position and its closest TCR residue in the murine dataset. The boundaries of the linear regression line indicate the 95% confidence interval

**Supplementary Figure 8.**
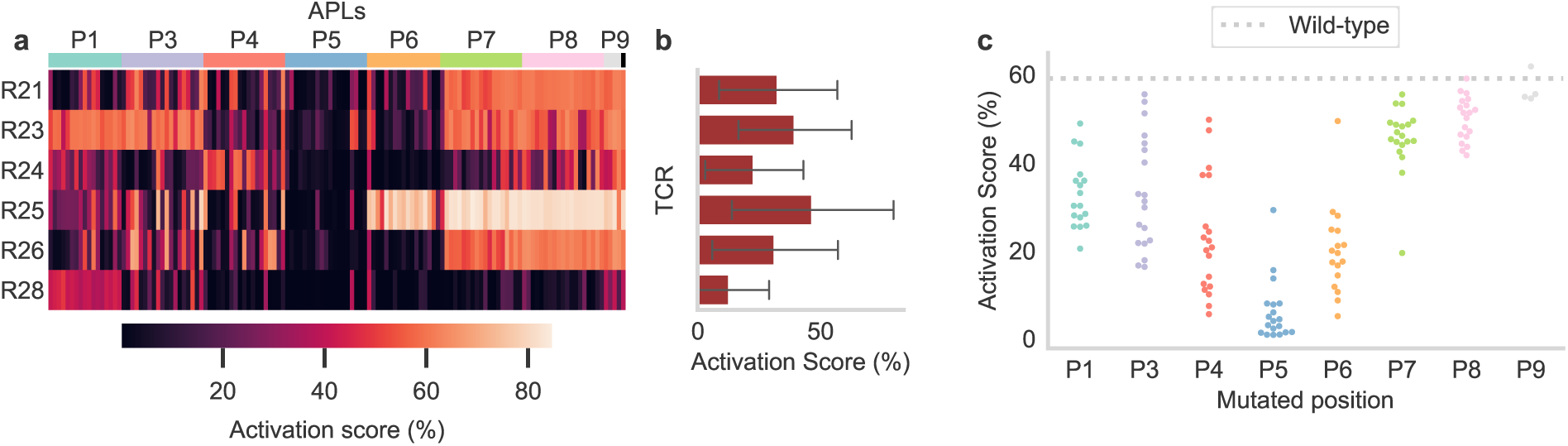
Unnormalized T cell activation for the TCRs and APLs of the human dataset. **a**, Unnormalized activation scores. **b**, Unnormalized activation scores averaged for all APLs per TCR. **c**, Unnormalized activation per APL over all TCRs.

**Supplementary Figure 9.**
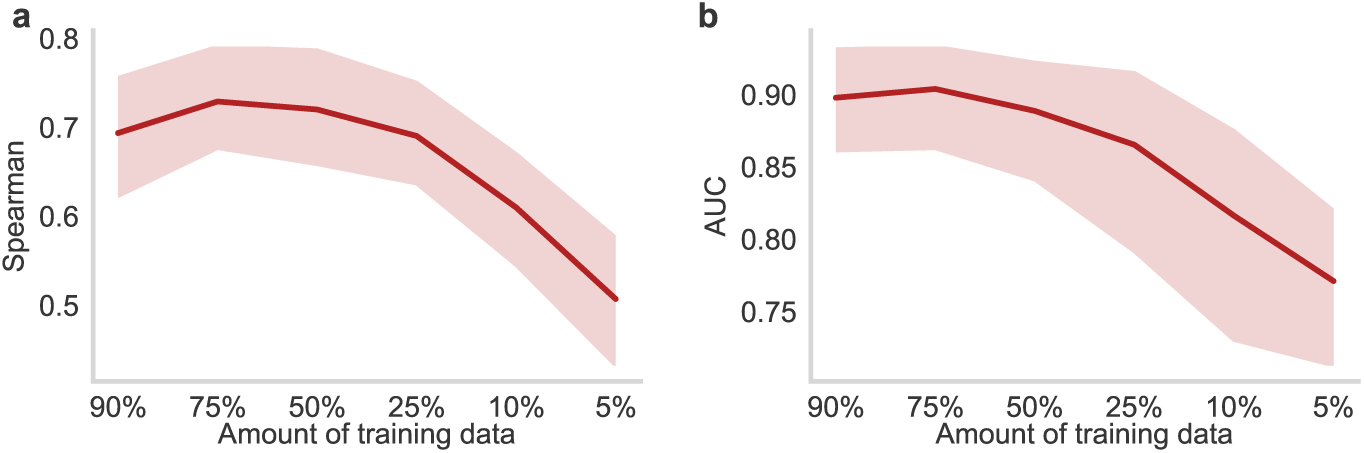
Performance on the human dataset on limited training data. Spearman correlation (**a**) and AUC (**b**) when a smaller amount of training data is used (average over ten repetitions with random subsets for each TCR). The boundaries of the mean line indicate the 95% confidence interval.

**Supplementary Figure 10.**
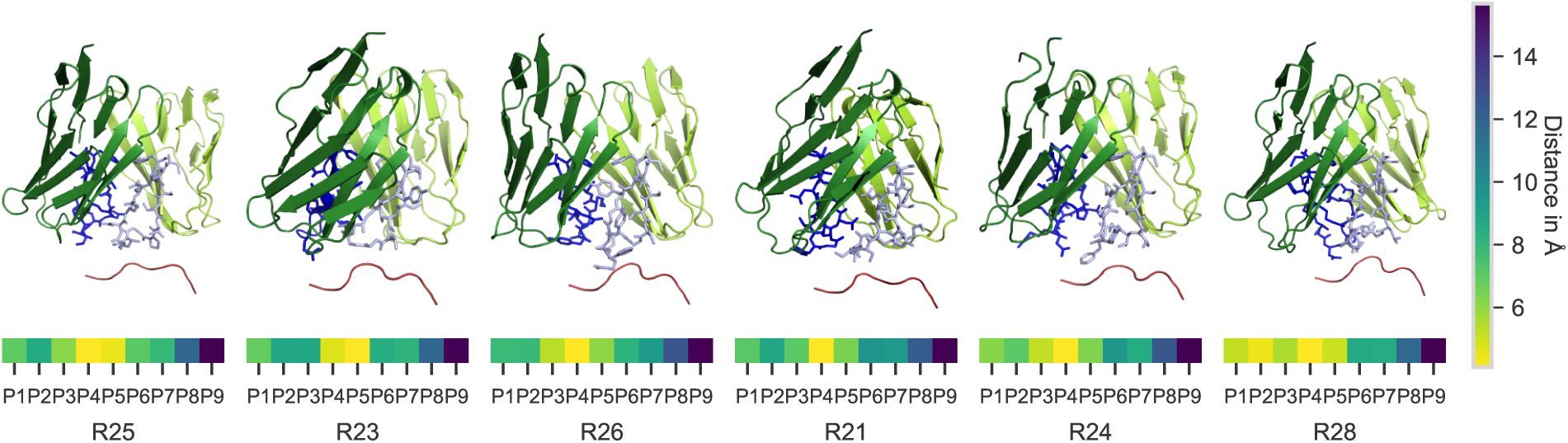
Structural Models of the human dataset. Predicted structures of the TCR and epitope, and minimal distance to the individual epitope positions for all receptors of the human datasets ordered by descending activation to the wildtype epitope.

**Supplementary Figure 11.**
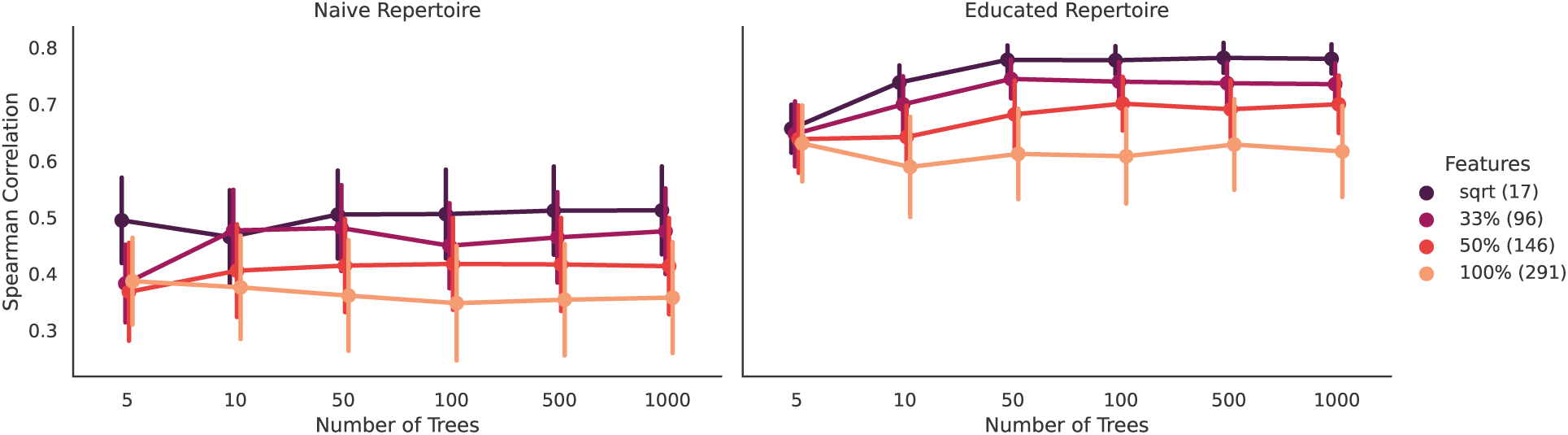
Performance of P-TEAM by number of trees in the random forest, and number of features used to build each tree. The Spearman correlation between the predicted and measured activation scores in a leave-TCR-out validation scheme is shown as a function of the number of trees used to train the random forests (*x*-axis), and the number of randomly-chosen features used to grow each tree.

## Supplementary Tables

**Supplementary Table 1.**
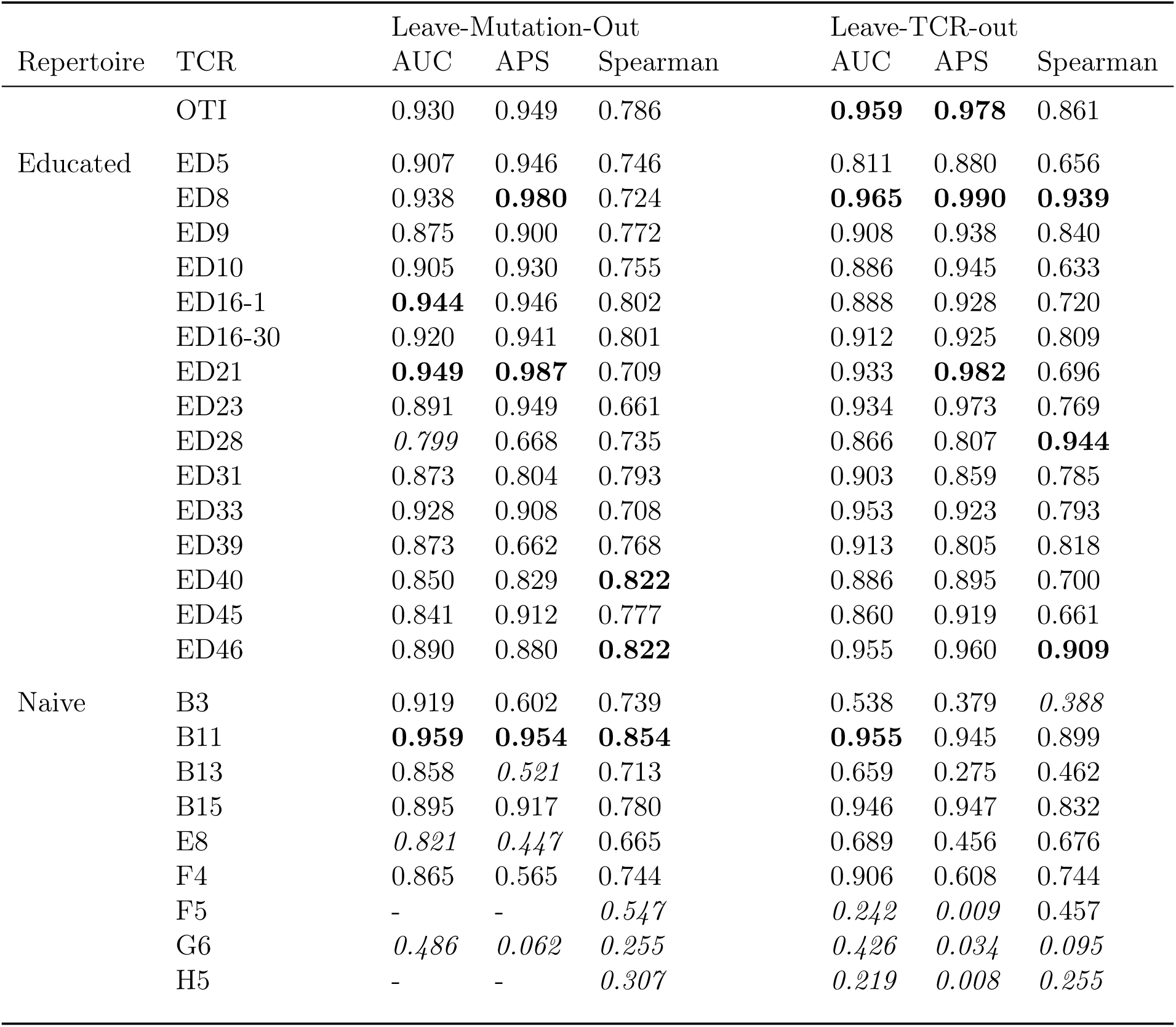
Performance of P-TEAM on the murine dataset. The classification performance is reported by the area under the receiver operator characteristic curve (AUC) and the average precision score (APS). The regression performance is reported by the Spearman’s rank coefficient. The three TCRs on which P-TEAM performed best and worst are shown in bold and italics, respectively, for each metric. AUC and APS of F5 and H5 could not be calculated as no sample was predicted as positive.

**Supplementary Table 2.** Performance of P-TEAM on the human dataset. The classification performance is reported by the area under the receiver operator characteristic curve (AUC) and the average precision score (APS). The regression performance is reported by the Spearman’s rank coefficient. The TCRs on which P-TEAM performed best and worst are shown in bold and italics, respectively, for each metric.

| TCR | Leave-Mutation-Out |  |  | Leave-TCR-out |  |  |
| --- | --- | --- | --- | --- | --- | --- |
|  | AUC | APS | Spearman | AUC | APS | Spearman |
| R21 | 0.906 | 0.882 | 0.682 | **0.938** | **0.916** | **0.800** |
| R23 | 0.892 | 0.872 | 0.744 | 0.788 | 0.842 | 0.708 |
| R24 | *0.761* | *0.322* | 0.700 | 0.618 | 0.275 | *0.427* |
| R25 | **0.949** | **0.933** | **0.846** | 0.809 | 0.864 | 0.681 |
| R26 | 0.920 | 0.901 | 0.780 | 0.916 | 0.899 | 0.766 |
| R28 | 0.911 | 0.791 | *0.654* | *0.561* | *0.183* | 0.509 |

## Supplementary Data

**Supplementary Data 1 | Educated repertoire of the murine dataset.** Unnormalized and normalized activation scores as well as sequences of the murine dataset for all 15 educated TCRs and OTI.

**Supplementary Data 2 | Naive repertoire of the murine dataset.** Unnormalized and normalized activation scores as well as sequences of the murine dataset for all 20 naive TCRs and two replicas of OTI (excluded from analysis).

**Supplementary Data 3 | Neo-antigen specific human dataset.** Unnormalized and normalized activation scores as well as sequences of the human dataset for all seven TCRs (R27 excluded from analysis).

**Supplementary Data 4 | Alligned TCR sequences.** *α*- and *β*-chain CDR3 sequences of the murine and human datasets aligned by the MUSCLE algorithm.

**Supplementary Data 5 | TCR distances.** Pairwise distances for the TCRs separated for the murine and the human dataset as defined by TCRdist.

**Supplementary Data 6 | Structural models of the murine dataset.** PDB files containing the structural models of the 36 murine TCRs, SIINFEKL, and the H2k^b^ allele.

**Supplementary Data 7 | Structural models of the human dataset.** PDB files containing the structural models of the 6 human TCRs, VPSVWRSSL, and the HLA-B*07:02 allele.

